# Visualizing the Transiently Populated Closed-State of Human HSP90 ATP Binding Domain

**DOI:** 10.1101/2022.03.16.484593

**Authors:** Faustine Henot, Elisa Rioual, Adrien Favier, Pavel Macek, Elodie Crublet, Pierre Josso, Bernhard Brutscher, Matthias Frech, Pierre Gans, Claire Loison, Jerome Boisbouvier

## Abstract

HSP90 are abundant molecular chaperones, assisting the folding of several hundred client proteins, including substrates involved in tumor growth or neurodegenerative diseases. A complex set of large ATP-driven structural changes occurs during HSP90 functional cycle. However, the existence of such structural rearrangements in apo HSP90 has remained unclear. Here, we identified a metastable excited state in the isolated HSP90 ATP binding domain. We used solution NMR and mutagenesis to characterize structures of both ground and excited states. We demonstrated that in solution the HSP90 ATP binding domain transiently samples a functionally relevant ATP-lid closed state, distant by more than 30 Å from the ground state. NMR relaxation and molecular dynamics were combined to characterize the energy landscape corresponding to the transition between these interconverting states. The precise description of the dynamics and structures sampled by human HSP90 ATP binding domain is a paramount piece of information for the future design of new therapeutic ligands.

## Introduction

Heat Shock Protein 90 (HSP90) is an ubiquitous ATP-dependant molecular chaperone involved in the folding of a plethora of client proteins, also modulating their cellular activities. HSP90’s activity is itself regulated by post-translational modifications such as phosphorylation, nitration, methylation or acetylation^1^ as well as a large number of different co-chaperones^2–7^, including Hop, PP5, p23, Sgt1, FKBP51/52, Cyp40, NudC and Cdc37. During its functional cycle, in order to fold client proteins, the chaperone HSP90 undergoes complex structural rearrangements driven by both ATP binding and hydrolysis^8,9^. Among the identified HSP90-client proteins, a large fraction is related to cancer such as steroid hormone receptors, tumor suppressor p53, telomerase, hypoxia-inducible factor 1α and kinases^10^. Furthermore, high cellular HSP90 expression levels are often associated with poor prognoses in many cancer types^11^. Therefore, human HSP90 has been identified as a major anti-cancer drug target. There are four HSP90 homologues in human cells: GRP94, TRAP1, HSP90α and HSP90β^12^. While GRP94 and TRAP1 are respectively found in endoplasmic reticulum^13^ or mitochondria^14^, respectively, both HSP90α and HSP90β isoforms are highly abundant in the cytoplasm, representing 1 to 2 % of cellular proteins^15–17^. Under stress conditions or in tumor cells, the level of expression of HSP90 increases up to 7% of the total amount of expressed proteins^10^.

HSP90 is a homodimeric chaperone of *ca*. 170 kDa. Each monomer is composed of 3 domains: C-terminal domain (CTD), middle domain, and N-terminal domain (NTD). The 13 kDa CTD is responsible for HSP90 dimerization, and harbours a MEEVD motif mediating the binding of various co-chaperones with tetratricopeptide repeat (TPR) domains^18–20^. The middle domain (*ca*. 40 kDa) is involved in binding of both client proteins and co-chaperones and, additionally, modulates hydrolysis of ATP bound to the NTD^3,21,22^. A long charged linker connects the middle domain and the NTD and modulates molecular interactions with client proteins^23^. Finally, the NTD (*ca*. 25 kDa) is implicated in client protein and co-chaperone binding^3^. HSP90-NTD possesses an ATP binding site with an unusual structure named Bergerat fold^24^. The particular environment of the

ATP binding site offers the possibility to develop HSP90-specific inhibitors that are not affecting the activity of most other ATP-binding proteins. ATP binding and hydrolysis in the NTD domain drive the structural rearrangements of HSP90 during its functional cycle^25–28^. Therefore, the HSP90 ATP binding site is the target of most therapeutic ligands developed so far, and more than 300 crystal structures of HSP90-NTD, bound to different ligands, are available in the PDB. A particular challenge for the design of new inhibitors against this important cancer target is the presence of a highly flexible ATP-lid segment that can cover the nucleotide/drug binding site in HSP90^29,30^.

In this article, we report on an atomic-resolution structural and dynamics investigation of HSP90-NTD conformations sampled in solution. We demonstrate that the ATP-lid segment of this crucial human chaperone, in addition to the well characterized open state, also transiently populates a closed conformation that requires structural rearrangement of peptide segments over a distance of up to 30 Å. Using solution NMR spectroscopy and molecular dynamics (MD) simulation, we could determine atomic-resolution models of the open- and closed-state conformations, as well as derive kinetic and thermodynamic information for this structural rearrangement occurring in solution. This is the first time that such an excited closed-state structure is reported for isolated apo HSP90-NTD. Our study reveals that the closure of the ATP-lid, observed during the ATP-driven functional cycle of HSP90, is already sampled in the human chaperone before binding of ATP. We anticipate that our results will be important for the future drug design of new inhibitors against this challenging drug target.

## Results

### Conformational variability of HSP90α-NTD

HSP90 has been identified as a major therapeutic target, especially against cancer^2,31,32^. Since the late 90’s a large number of competitive inhibitors, targeting the ATP binding site of the HSP90α-NTD has entered clinical trials^33–35^. In the context of these drug development efforts, more than 300 atomic resolution structures of isolated HSP90*α*-NTD in presence of various ligands have been determined, using X-ray crystallography. These structures can be separated into 8 main groups (Fig. 1a-c) showing distinct conformational properties. Pairwise superimposition of the HSP90α-NTD centroid structures representing each cluster reveals that the main structural differences are located on the segment covering the nucleotide/drug binding site. While large parts of these 8 centroids superimpose with an average root-mean-square deviation (RMSD) of *c.a.* 0.2 Å, the ATP-lid^29^ (defined here from residues M98 to V136 including three helical segments) shows higher heterogeneity with an average pairwise RMSD of 1.4 Å (Fig 1d). In all the apo and nucleotide- or ligand-bound structures of the isolated HSP90*α*-NTD, the ATP-lid is in the so-called open state^27^ and does not cover the ATPase site and corresponding drug binding site. The observed structural variability of HSP90α-NTD structures indicates that the ATP-lid, in the open state, can adopt various conformations in a crystalline environment depending on the crystallization conditions, and the stabilization by ligands or additives. However, a detailed picture of the conformational properties of the ATP-lid segment in solution is of utmost importance for future drug design, as this segment is located next to the major drug binding site of human HSP90. Therefore, we initiated a structural and dynamic investigation of HSP90-NTD using solution NMR spectroscopy to characterize at atomic resolution the conformations sampled by the ATP-lid.

**Figure 1:**
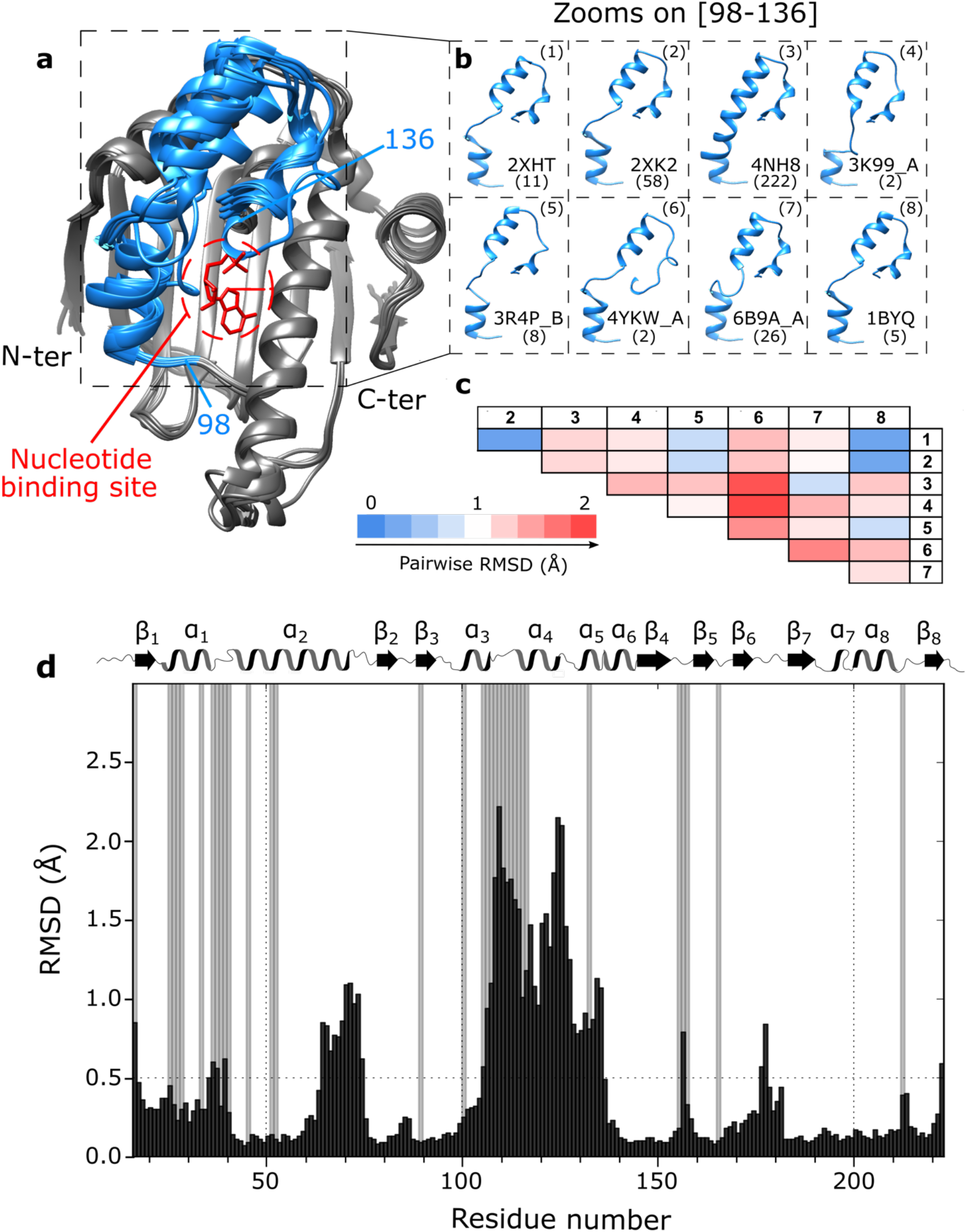
Analysis of available structures of human HSP90*α*-NTD. **a)** Superimposition of 8 centroids representing the 8 clusters describing the 334 structures of isolated HSP90*α*-NTD available in the Protein Data Bank (on January the 5^th^ of 2021). Clustering was performed using the MaxCluster program (http://www.sbg.bio.ic.ac.uk/maxcluster) with an Average Linkage type of hierarchical clustering and a threshold value of 1.05. In blue is depicted the segment [98-136] and in red the nucleotide. **b)** Zooms on the segment [98-136] for all the 8 centroids superimposed in a). PDB ID and number of structures present in each cluster are disclosed next to each centroid. **c)** Table representing pairwise RMSD in Å between Cα backbone atoms of each pair of the 8 centroids. (1: 2XHT, 2: 2XK2, 3: 4NH8, 4: 3K99_A, 5: 3R4P_B, 6: 4YKW_A, 7: 6B9A_A, 8: 1BYQ). Going from blue to red the RMSD values increase. d) Histogram of the averaged pairwise RMSD between Cα backbone atoms of the 8 centroids in Å as a function of the residue number (black). Grey bars represent non-assigned backbone residues^38^. On top of the histogram: secondary structure elements as a function of the residue number (*α* for helices and *β* for sheets).

### The ATP-lid of HSP90-NTD populates two distinct conformations in solution

Previous NMR investigations reported that human HSP90*α*-NTD amide backbone NMR signals from N105 to K116 in the ATP-lid segment were undetectable^36–38^ (Fig 1d). This observation suggests that the ATP-lid is undergoing conformational exchange on the micro- to millisecond time scale. The broadening of backbone NMR signals in the ATP-lid is also particularly detrimental for NMR-distance restraint based structure determination using uniformly ^13^C,^15^N-labeled HSP90*α*-NTD sample. In order to overcome this problem, we decided to produce perdeuterated HSP90α-NTD, specifically ^13^CH_3_-labeled on the A*^β^*, I*^δ^*^1^, L*^δ^*^2^,M*^ε^*,T*^γ^* and V*^γ^*^2^ methyl positions^39^. The resulting 87^1^H,^13^C-labeled methyl groups all gave rise to detectable NMR signals. Most interestingly, 5 of these methyl groups (L107, T109, I110, A111 and T115) belong to the previously unassigned stretch of residues in the ATP-lid (Supporting Fig. S1). Unambiguous sequence-specific ^1^H and ^13^C resonance assignments of all detected methyl groups was obtained from a set of through-bond correlation experiments^38^, aided by an extensive mutagenesis-driven assignment of 56 key methyl probes^40^ (Supporting Fig. S2).

Next, we recorded methyl NOESY spectra to derive a set of inter-methyl distance restraints. The applied labeling scheme that yields protonation only on the methyl moieties at the extremity of hydrophobic side chains is particularly useful to extract long-range distance restraints between methyl groups that are separated by up to 10 Å^41^. For HSP90*α*-NTD, 597 inter-methyl NOEs (Supporting Fig S3A and Table S1) could be detected, from which 111 distance restraints were derived that involve at least one methyl group belonging to the ATP-lid segment. All these structurally meaningful restraints were assigned exclusively based on chemical shifts. The inter-methyl distance restraints were complemented by 41 short-range backbone H_N_-H_N_ distance restraints (Supporting Fig S2A and Table S1) derived from a 3D ^15^N-edited NOESY spectrum, and 54 backbone φ,ψ-dihedral restraints derived from characteristic backbone chemical shifts^38,42^.

Based on this set of experimentally derived structural restraints, we have performed structure calculations using a simulated annealing protocol in torsion angle space^43^ to determine the structure and position of the ATP-lid segment relative the rigid HSP90*α*-NTD scaffold comprised by the two segments [11-97] and [137-223]). To our surprise, the calculated structures showed a large number of distance violations (25) and a high final target energy (74 kJ.mol^-1^). Careful analysis of the reported distance violations revealed a number of NMR-derived restraints that are incompatible with a single structure (Fig. 1a). As an example, a pair of inter-methyl NOEs is detected between the methyl group of I131-δ_1_ located in the ATP-lid and the methyl moiety of L29-δ_2_ in helix*−*1, while another set of NOE correlation peaks indicate that the preceding residue M130-*ε* in the ATP-lid is spatially close to L64δ_2_ and T65-*γ*, two methyl groups located at the end of the long helix-2 (Fig. 2ab). However, according to the previously determined X-ray structures of HSP90-NTD (Fig. 1), L29 is more than 30 Å away from L64 and T65. The specific assignment of these methyl groups, the analysis of the through-bond sequence specific assignments of backbone nuclei and the transfer to methyl-moieties were double checked^38^ and completed by an extensive mutagenesis driven assignment of 56 keys methyl probes^40^ (Supporting Fig. S2) allowing us to exclude misassignment of these NOEs. As the NMR-derived distance restraints could not be satisfied with a single structure, we made the hypothesis that the ATP-lid populates several conformations. Therefore, an additional round of structure calculation was performed^44^ assuming two co-existing conformations of the ATP-lid segment (Fig. 2c), and allowing each experimental distance restraint to be satisfied by either one of the two structures, or both. The introduction of two distinct conformational states in our calculation protocol allowed to satisfy all experimental restraints, and also decreased the final target energy (Fig. 2d) by a factor of 16. Additional calculations including a third state did not result in further improvement of the target energy (Fig. 2d), indicating that a model including two states is sufficient to satisfy all experimental data.

**Figure 2:**
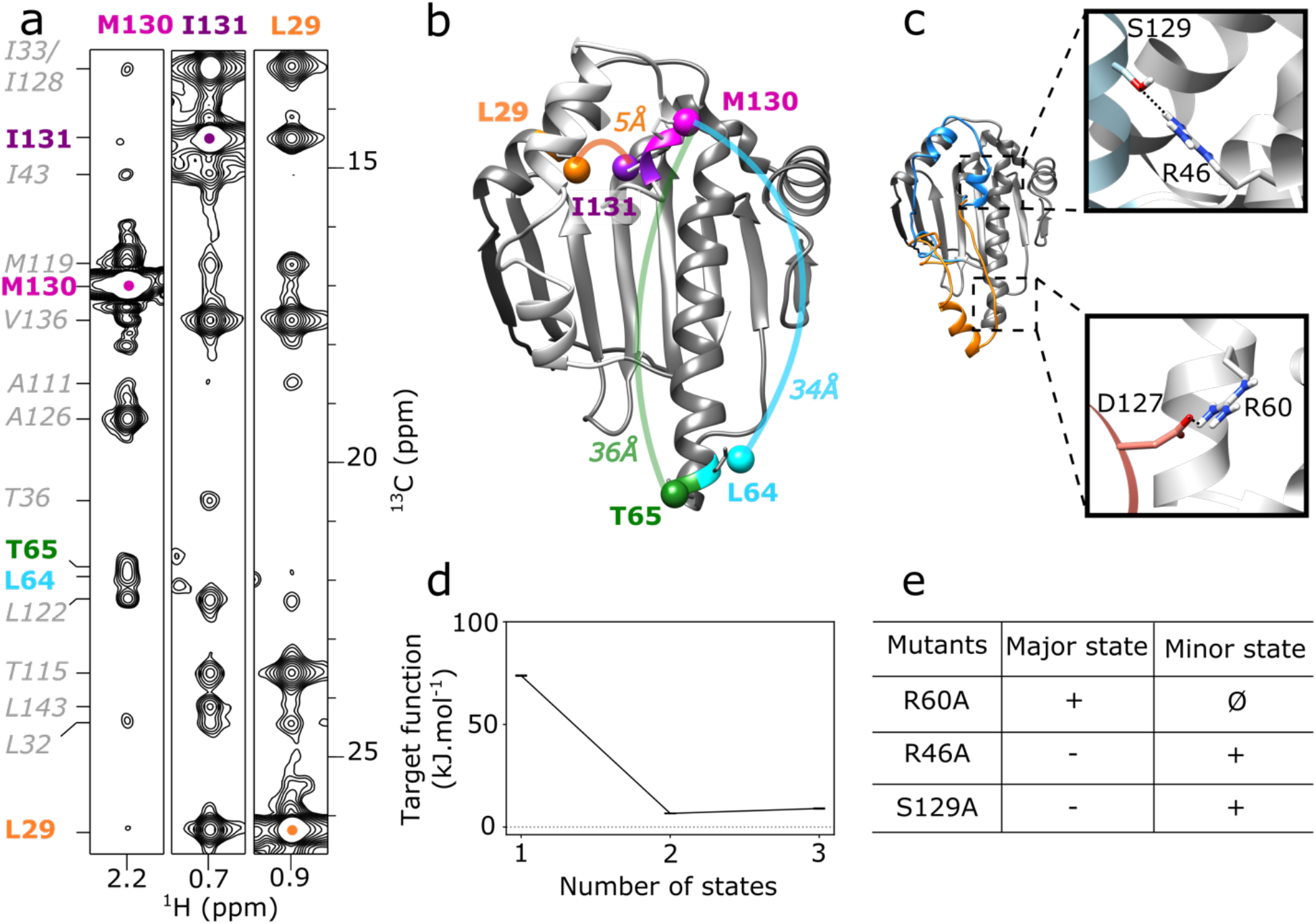
HSP90*α* NTD samples two states in solution. **a)** 2D-strips extracted from ^13^CH_3_-edited 3D NOESY, at ^13^C frequencies of M130, I131 and L29 methyl groups of WT-HSP90α-NTD. The 3D HMQC-NOESY-HMQC experiment was acquired at 25°C on a spectrometer operating at a ^1^H frequency of 950 MHz. **b)** Representation on a 3D structure of HSP90α-NTD (PDB code: 1YES) of examples of intermethyl NOEs involving M130 and I131 with methyl groups distant by more than 30 Å. **c)** Representatives two states of HSP90α-NTD calculated simultaneously^54^ to satisfy all NMR restraints acquired on WT-HSP90α-NTD. Boxes correspond to zooms on structure displaying interactions stabilizing the conformation of the ATP-lid either in closed (orange) or open conformation (blue) **d)** Variation of average CYANA target function for the 20 best structures according to the number of states used to calculate each conformer**. e)** Table summarizing intensity increases (+), decreases (-) or disappearance (Ø) of characteristic NOEs corresponding to HSP90α-NTD open and closed states (Supporting Table S2).

When the two final average structures are superimposed on the common peptide segments [11-97] and [137-223], the backbone RMSD of the ATP-lid segment is *ca.* 20 Å (Fig. 1c). The first state corresponds to the structure with the ATP-lid in an open position, similar to the bundle of representative structures previously resolved for the isolated HSP90*a*-NTD (Fig. 1a). The second state shows the ATP-lid in a closed conformation, covering the entire ATP-binding site, and helix*−*4 located near the end of long helix*−*2. A careful analysis of these structures showed that 22 NOE-derived distance restraints are satisfied only by the ATP-lid open state, while 5 distance restraints are specific to the ATP-lid closed state (Fig. 2c, Supporting Table S2), and the remaining 125 restraints are satisfied in both states.

### Stabilization of ATP-lid closed and open states by mutagenesis

Our structural model of the ATP-lid closed state points toward a salt bridge involving residues R60 and D127 as a potential contributor to the enthalpic stabilization of this so far unknown conformation of HSP90*α*-NTD (Fig 2c). Therefore, we speculated that HSP90*α*-NTD mutants suppressing the R60/D127 electrostatic interaction would push the population equilibrium toward the ATP-lid open conformation. Two ^13^CH_3_-labeled samples of single-point HSP90*α*-NTD mutants (D127A and R60A) were prepared and analyzed by NMR. While the D127A mutant showed NMR spectral signatures characteristic of a structurally heterogeneous sample, the R60A mutant yielded a single set of ^1^H-^13^C correlation peaks -similar to the WT (Supporting Fig. S4a). Analysis of a ^13^CH_3_-edited NOESY spectrum recorded for this R60A mutant revealed that all expected NOE correlation peaks specific of the ATP-lid closed state were no longer detected for this mutant (Supporting Fig. S5). This result indicates that disruption of the R60/D127 salt bridge decreases the population of the ATP-lid closed state to a level that is no longer detected by NOESY-type NMR experiments.

In a similar way, our structural model of the ATP-lid open state allowed us to identify a stabilizing hydrogen bond between the side chains of R46 and S129 (Fig 2c). Again, we produced ^13^CH_3_-labeled samples of single-point HSP90*α*-NTD mutants (R46A and S129A), in order to destabilize the open state and increase the population of the closed state. For both mutants, homogeneous ^1^H-^13^C correlation spectra were obtained (Supporting Fig. S4b), and methyl NOEs characteristic of both the open and closed states were detected. Still, these mutations resulted in a shift of population from the open to closed state, as the peak intensities of NOE correlation peaks specific for the closed state were enhanced by about a factor *ca.* 2.8 with respect to those characteristic of the open conformation (Supporting Fig. S5). The observation of population shifts induced by single-point mutations derived from our structural models of the ATP-lid closed and open states also strongly supports our 2-states model and the existence of an ATP-lid closed state in WT-HSP90*α*-NTD in solution.

### NMR refinement of structural models for the ATP-lid open and closed states

Since the HSP90*α*-NTD mutants possess populations of the two ATP-lid conformations strongly skewed toward either an open or a closed state, they provide an opportunity to further refine structural models. For structure calculation of the ATP-lid open state, a set of 114 distance restraints, involving at least one methyl group of the ATP-lid segment, was derived from the 3D ^13^CH_3_-edited NOESY spectrum acquired using ^13^CH_3_-labeled sample of R60A-HSP90*α*-NTD. As the experimental set of distance restraints is self-consistent, calculations were performed assuming a single ATP-lid open state for each calculated conformer. No violation of experimental restraints was observed in the 20 best calculated structures. After a final refinement step using molecular dynamics in explicit water and in presence of experimental restraints^45^, the structure of the ATP-lid segment can be superimposed on backbone atoms with a RMSD to the average structure of 0.9 Å (Fig. 3a, Table S3). The calculated ATP-lid open conformation forms three helices ([100-104], [115-123] and [128-135]) separated by short loop segments. As the backbone signals were not detectable for the segment [105-115], this part of the ATP-lid is less well defined due to a limited number restraints.

**Figure 3:**
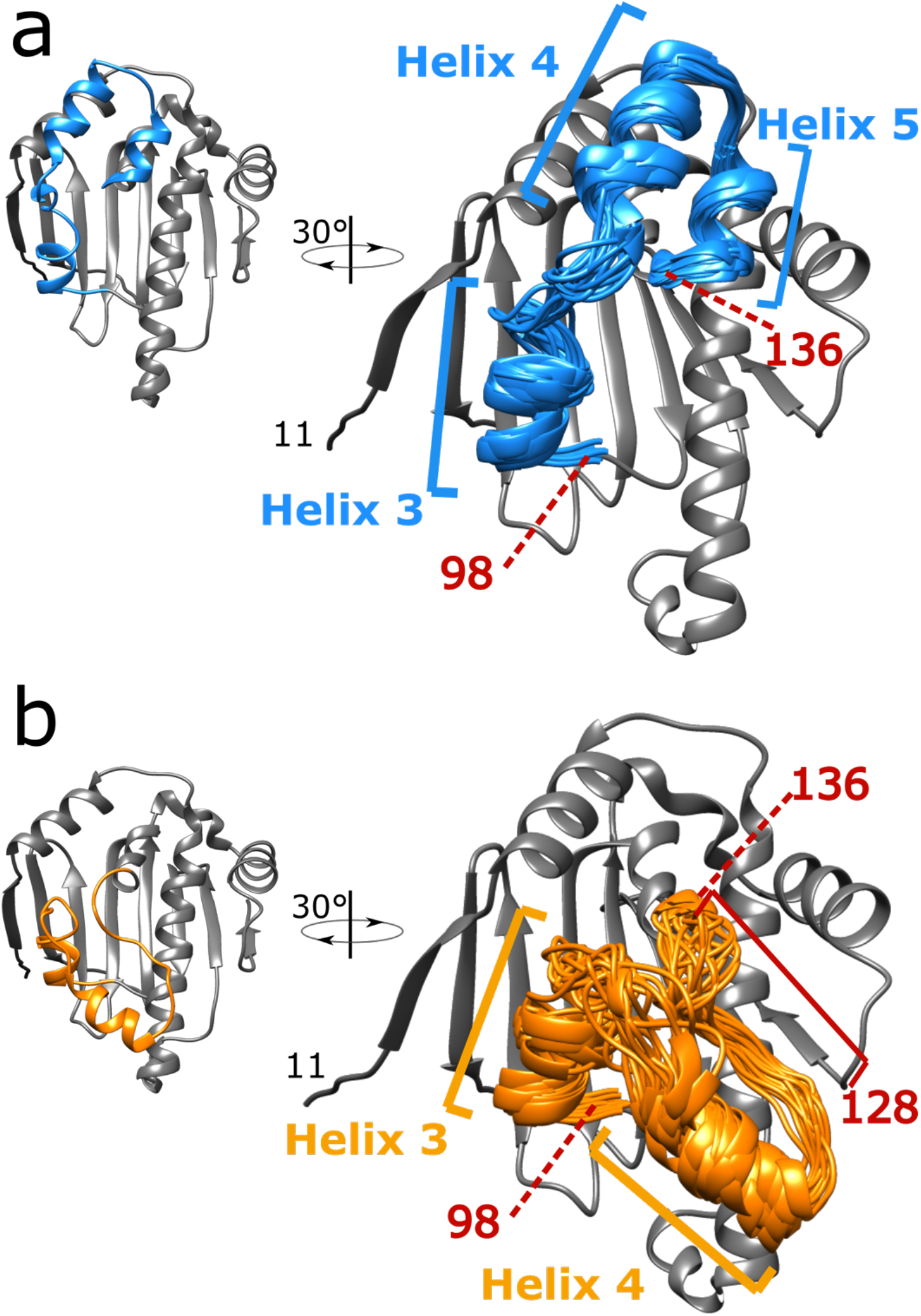
Solution structure ensembles of HSP90α-NTD ATP-lid open and closed states. For each state, the 20 best CYANA conformers were selected for further restrained molecular dynamics refinement in explicit water. For each panel, the centroid representative conformer of the ensemble is presented on the left in an orientation similar to Fig 2b,c. On the right the structure ensemble is tilted by 30° and was superimposed on the coordinate of the centroid conformers. In **a)** structure ensemble for ATP-lid open/ground-state calculated using NMR structural distance restraints obtained using R60A-HSP90α-NTD sample. In **b)** Structure ensemble for ATP-lid closed/excited-state calculated using NMR structural distance restraints obtained using R46A-HSP90α-NTD sample. Helices 3, 4 and 5 correspond to ATP-lid helices. The positions 98 and 136 of the pivots are also indicated. The location of the unfolded *a*_5_ helix in the closed state is also indicated (between 128 and 136).

For refinement of the ATP-lid closed state we based our structure calculation on NOE-derived distance restraints from the R46A mutant, as this mutation is located far away from ATP-lid in closed state contrary to S129A mutant which may affect ATP-lid structure. For R46A mutant NOEs that are specific to the ATP-lid open state can still be detected, although at reduced intensity, we used the refined structure of the ATP-lid open state to identify and filter-out restraints specific to this conformation (Supporting Table S2). The remaining 81 inter-methyl NOEs, completed with 36 dihedral- and 41 backbone distance restraints (Supporting Table S1), were used to refine the structure of the ATP-lid segment in the closed state on the rigid HSP90*α*-NTD scaffold. The final structural ensemble, superimposable to the average structure with a backbone RMSD of 1.6 Å (Fig. 3b), did not show any violation of experimental restraints. The hinge residues allowing the ATP-lid to switch from an open to a closed conformation are residues [109-111] and [135-136]. While the helix-3 and helix-4 are conserved in this structure, the segment corresponding to helix*−*5 in the ATP-lid open state is in an extended conformation in the closed state, thus enabling formation of a salt-bridge between R60 and D127 as well as methyl-methyl contacts between ATP-lid A121, A124, A126, M130 side-chains and methyl moieties L64 and T65.

### The ATP-lid closed state is a metastable excited state

To study the stability of both HSP90 ATP-lid open and closed states, we have explored the conformational landscape around these two families of conformers using molecular dynamics simulations (without experimental restraints). Representative conformers of the structural ensembles obtained for the open and closed states were used as starting models for forty 1 μs-molecular dynamics simulations. Simulations starting from an ATP-lid in the open state remained stable over the whole 1-μs trajectory. More quantitatively, the pair-wise C*α*-RMSD over 2000 conformations extracted every 10 ns from the twenty 1-μs long trajectories remains low (Fig 4.a bottom left corner), with an average of 4.0 Å. Furthermore, these structures satisfy the NOE contacts characteristic of the ATP-lid in the open state (Fig. 4.b), even if no restraints were applied during the molecular dynamics simulations. In contrast, the 2000 conformations extracted from the 20 trajectories starting with the ATP-lid in the closed state showed a larger averaged pairwise RMSD (9.3 Å, see the top right corner of Fig 4.a). This structural diversity may originate from a complex energy landscape around an excited state, and/or from the uncertainties of the initial experimental structures. Roughly, the trajectories show three typical behaviors. The first case scenario is represented by 7 trajectories (among 20) that were considered as stable, *i.e.,* they explore conformational space around the starting conformation without large violations of the NOE contacts that are characteristic of ATP-lid in the closed state during the whole 1 μs-simulations (Fig. 4.d-I and Supporting Figure S6). This indicates that the refinement procedure has provided a state that is metastable. In contrast, among the unstable trajectories, about 9 show a transition toward conformations almost compatible with the NOE restraints characteristic of the open state (Fig. 4.d-II). They are characterized by RMSD values, in the anti-diagonal corners of Fig 4.a, as low as 1.5 Å. Finally, in 20 % of the trajectories, the ATP-lid populates conformations that are neither clearly the closed, nor the open state (Fig. 4.d-III). The observation of several spontaneous transitions of the ATP-lid from the closed to the open state, and the stability of the MD trajectories starting from the open state, indicate that the ATP-lid open state (the only state observed by X-ray crystallography - Fig. 1), is the ground state of HSP90-NTD, while the ATP-lid closed state is a metastable, less populated excited state. This conclusion is also supported by the observation of a small number (5) of characteristic NOEs for the closed ATP-lid state compared to a larger number (22) detected for the open ATP lid ground state (Supporting Table S2).

**Figure 4:**
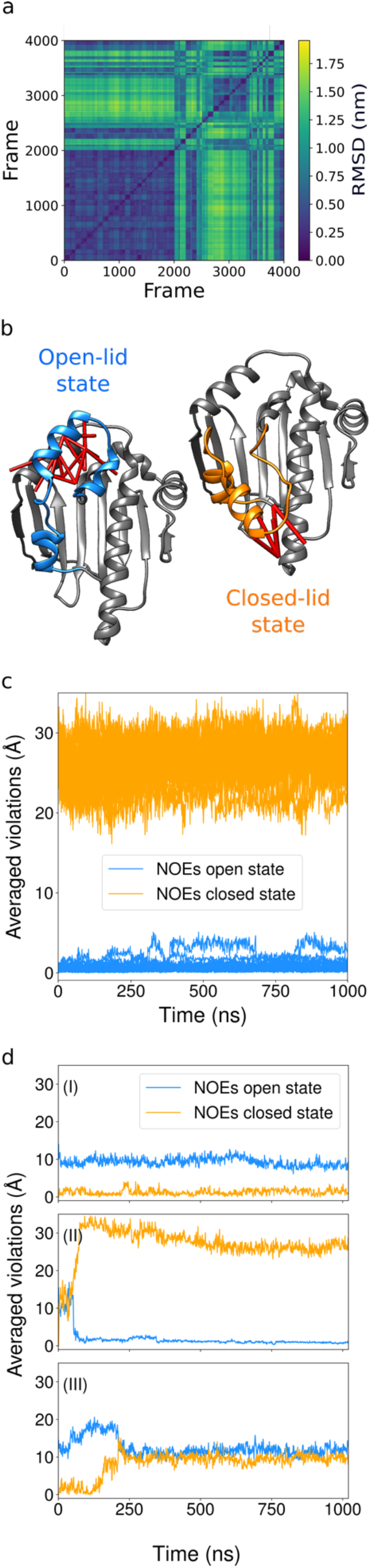
Molecular dynamic investigation of HSP90α-NTD ATP-lid open and closed states. **a)** Pairwise RMSD (in nm) among the ATP-lid structures obtained in the forty Molecular Dynamics simulations of 1 µs duration each. The initial models are the ensemble of experimentally refined structures (Fig. 3). Hundred frames, separated by 10 ns from each other, are extracted from each simulation. The first 2000 frames are extracted from simulations starting with ATP-lid open-state conformers. The last ones are extracted from simulations starting with ATP-lid closed-state conformers. Going from blue to yellow, the RMSD values increase. **b)** Representation of NOE distance restraints characteristics of ATP-lid open and closed states on the corresponding structure. **c)** Violations of characteristics NOEs monitored during the molecular dynamics simulations performed without restraints, for the 20 simulations starting with the ATP-lid in the open state (See online methods for definition of computed violations). The values are averaged over all the ATP-lid open (blue) or closed (orange) states specific distances restraints (Supporting Table S2). **d)** As (c), for a typical simulation starting with the ATP-lid in the closed state that remains stable (I), undergoes a transition towards the ATP-lid in the open-state (II), or derives towards region of conformational space that is neither the closed, nor the open ATP-lid state (III).

### ATP-lid open and closed states exchange on the millisecond time scale

The fact that a single set of NMR signals is detected for HSP90*α*-NTD (WT, R60A and R46A mutants) indicates that the ATP-lid open and closed states are in fast exchange with respect to the NMR time scale (*τ*_ex_ < ∼10 ms). The observed severe line broadening of amide backbone resonances in the ATP-lid segment also points toward exchange dynamics on the micro- to millisecond time scale. In order to further quantify the kinetics (and thermodynamics) of the ATP-lid structural rearrangement, NMR relaxation-dispersion experiments were performed at 2 different magnetic field strengths using ^13^C-^1^H methyl, as well as ^15^N backbone amide probes. These data reveal that conformational exchange is mainly sensed by nuclei in the ATP-lid segment, as well as protein regions that are in close contact with the ATP-lid (Fig. 5a). Particularly strong exchange-contributions were detected in the hinge regions of the ATP-lid segment: residues [T109-T115] and V136. Assuming a simple two states model to describe the structural rearrangement of the ATP-lid, a systematic grid search of the model parameters fitting the relaxation dispersion data detected on backbone and methyl probes was performed. A single minimum was identified, indicating that the experimental data can be interpreted with the presence of a single low populated minor state (Supporting Fig. S7). A global fit of the ^13^CH_3_-CPMG relaxation dispersion, characterized by a higher signal to noise ratio, enabled us to determine the global exchange rate k_ex_=k_1_+k_-1_= 2490 ± 61 s^-1^ and relative populations of the 2 states of 96.8 ± 0.1 % (ground state: ATP-lid open state) and 3.2 ± 0.1 % (excited state: ATP-lid closed state). This corresponds to half-life times of 8.6 ± 0.4 ms for the ATP-lid open state, and 0.29 ± 0.01 ms for the ATP-lid closed state. These populations and exchange rates correspond to a difference of Gibbs free energy of 8.3 ± 0.2 kJ.mol^-1^ between the 2 states, with an activation energy (from the ground to the transition state) of 61.0 ± 0.2 kJ.mol^-1^ ^46^.

**Figure 5:**
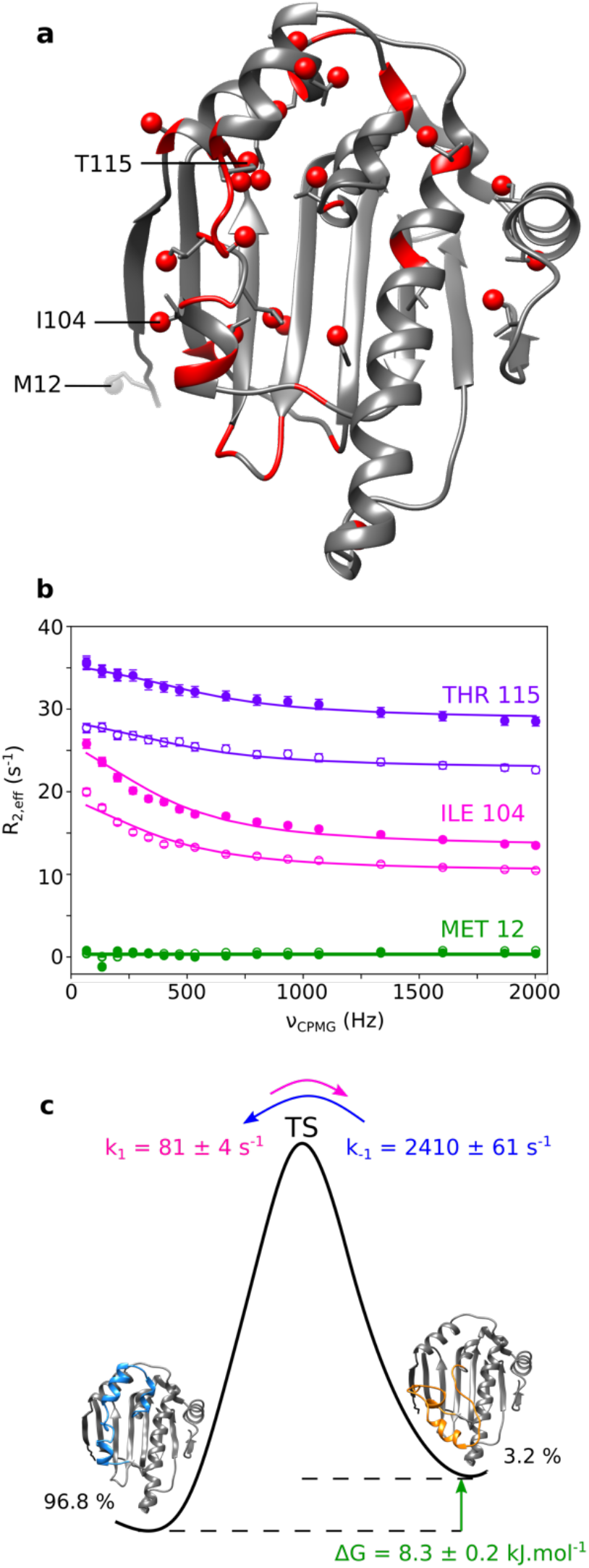
Relaxation dispersion study of HSP90α-NTD. **a)** 3D structure of human HSP90α-NTD displaying in red methyl- and backbone ^15^N-probes for which conformational exchange in the μs-ms time scale was detected. **b)** Examples of ^13^CH_3_ MQ CPMG relaxation dispersion profiles of Thr-115, Ile-104 and Met-12 plotted in purple, red and green, respectively. The filled circles for each color represent data acquired at 850 MHz and the empty circles represent data acquired at 700 MHz. Data displayed were acquired at 293 K using a U-[^2^H, ^15^N, ^12^C], Ala-[^13^C^1^H_3_]^β^, Ile- [^13^C^1^H_3_]^δ1^, Leu-[^13^C^1^H_3_]^δ2^, Met-[^13^C^1^H_3_]^ε^, Thr-[^13^C^1^H_3_]^γ^, Val-[^13^C^1^H_3_]^γ2^ HSP90-NTD sample. Experimental data were fitted to a two-sites exchange model (global fit of 21 relaxation dispersion curves). Precision on the data was estimated using Monte Carlo. **c)** Schematic diagram of the energy landscape for the exchange between the ATP-lid open (ground) and closed (excited) states of HSP90α-NTD. Both thermodynamics and kinetics parameters of the exchange, extracted at 293 K, are displayed. Activation energies can be estimated using Eyring equation (assuming κ =1). From the ground state to the transition state: ΔG* = 61.0 ± 0.2 kJ.mol^-1^ and from the excited state to the transition state: ΔG* = 52.7 ± 0.1 kJ.mol^-1^.

## Discussion

In this work, we have demonstrated that the ATP-lid segment of human HSP90*α*-NTD samples two distant conformations in solution. We have solved the solution structures of this important anti-cancer drug target with the ATP-lid either in an open or closed conformation. Such a closed state with the ATP-lid covering the nucleotide/drug binding site has never been observed before for isolated HSP90*α*-NTD, neither in the apo state nor in complex with nucleotides or drugs (Fig. 1). In addition to our NMR data, we have used mutagenesis and molecular dynamics simulation to independently validate the existence of this meta-stable, low populated closed state in human HSP90*α*-NTD. The calculation of Evolutionary Couplings (ECs) from sequence alignment of multiple homologous proteins is an emerging approach to identify conserved contacts and complement conventional approaches of protein structure determination^47^. Starting from an alignment of 4414 homologous sequences of HSP90-NTD, 71 highly probable ECs (p > 0.85) involving at least one residue of the ATP-lid segment (Supporting Fig. S8) were computed^47^. Interestingly, we found that both, the open and closed conformations are necessary to account for the structural contacts revealed by these ECs (Supporting Fig. S8), further supporting the existence of an ATP-lid closed state.

Using Molecular Dynamics simulations and NMR relaxation-dispersion experiments, we could establish that the ATP-lid in the open conformation is thermodynamically more stable than the ATP-lid in closed state, and that the population of the exited (closed) state is only 3-4 % at room temperature (Fig. 5). The open and closed states interconvert at a rate (k_ex_ = k_1_ + k_-1_) of 2.5 kHz, which seems to be very fast for such an extensive structural rearrangement. However, structural rearrangements at fast time scales were already reported for other conformational exchange events in proteins. An example is EIN, a 128 kDa homodimeric protein that switches from an open to a closed state at a speed faster than 10 kHz^48^. Furthermore, PET fluorescence quenching experiments by Schulze *et al.*^30^ monitoring yeast HSP90-NTD ATP-lid structural rearrangement in isolated monomeric constructs reveals a motion of the ATP-lid with a characteristic rate of 1.5 kHz, even though a full closure of the lid over the nucleotide/drug binding site has not been observed. This frequency range is similar to our results obtained by NMR relaxation dispersion experiments for human HSP90*a*-NTD. The transition pathway between open and closed ATP-lid states, distant by up to 30 Å, is obviously more complicated than a simple two-state model used here for the quantitative analysis of our NMR data. A more detailed analysis of the computed MD trajectories allows to obtain further information on the kinetics of the structural changes observed for the open-closed transition. The most pronounced rearrangements between the open and closed states are the reorientation of helix-3 and helix-4, as well as the folding/unfolding of helix*-*5. The MD trajectories describing a full transition from the closed to the open state show that the ATP-lid can tilt very rapidly, typically in a few hundreds of ns, while the folding of helix-5 and the associated local restructuration were not systematically observed during the 1-μs simulations (Supporting Figure S9). Therefore, we may conclude that the transition pathway involves several structural rearrangements occurring at different timescales, with the unwinding/folding of the helix-5 being one of the rate limiting steps of the transition between ATP-lid open and closed states.

To assist the folding of client proteins, the ATPase HSP90 interacts with a large number of co-chaperones and undergoes a conformational cycle in which a complex set of structural changes occurs. In currently proposed models of the HSP90 functional cycle, the two N-terminal domains of HSP90 transiently dimerize upon binding of ATP, client proteins and co-chaperones^49^. Dimerization is associated with an exchange of the first *β*-strand (β1) between both N-terminal domains, and a switch of the ATP-lid from an open to a closed state^29,30,50,51^. Cryo-EM structures of full-length HSP90 co-vitrified with ATP, a co-chaperone and a client protein^52,53^ reveal an ATP-lid segment in a closed conformation associated to a dimerization of the N-terminal domains and a swap of the β1-strands between both N-terminal domains (Fig. 6a-c). To investigate whether the closed ATP-lid conformation is responsible for dimerization of HSP90-NTD, we have identified 16 methyl-methyl proximities in the N-terminal domain of full-length human HSP90α (PDB: 7L7J) that are characteristic of the dimeric HSP90 with a closed ATP-lid and swapped β1-strands (Supporting Table S4). However, investigation of the inter-methyl NOEs observed for the HSP90-NTD mutant R46A, showing an enhanced population of the ATP-lid closed-state ensemble, revealed the absence of all the putative NOEs characteristic of a dimerization of HSP90α-NTDs, confirming that the closed state of HSP90α-NTD stays monomeric in solution.

**Figure 6:**
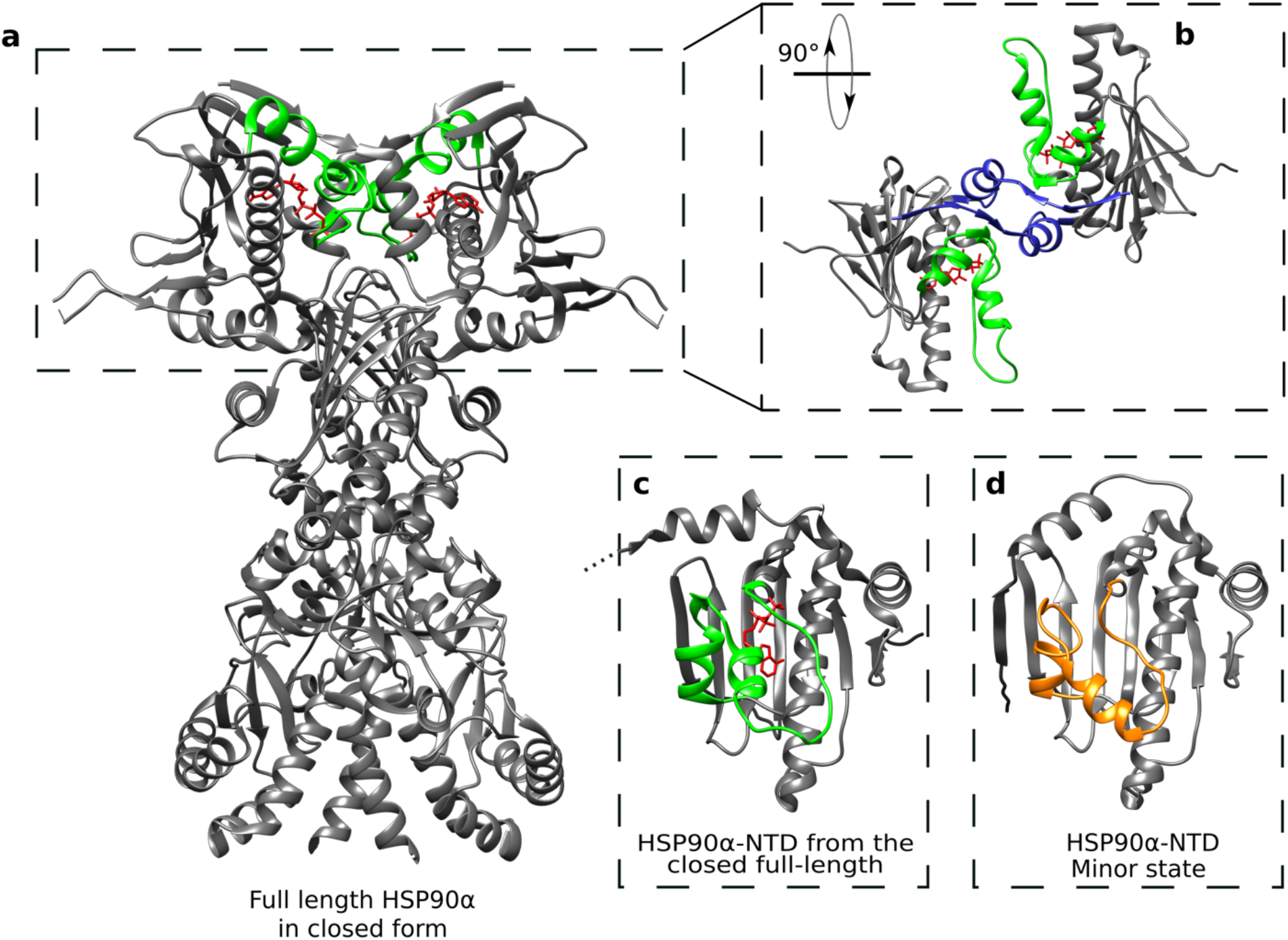
Structure comparison of HSP90α-NTD excited state with full-length dimeric HSP90 α functional cycle intermediate. **a)** Structure of the homodimer full-length HSP90α in closed form stabilized by p23, FKBP51 and ATP (PDB: 7L7J^53^). The segment covering the nucleotide binding site and ATP are colored in green and red, respectively. **b)** Zoom on the two N-terminal domains of the full-length HSP90α, turned by 90°. The blue segments represent the β strands exchanging between the two chains of the homodimer. **c)** N-terminal domain of HSP90α (PDB: 7L7J). d) Average structure of the calculated ensemble for ATP-lid closed (excited) state of apo HSP90α-NTD. The segment 98-136 is represented in orange.

The NMR-derived ATP-lid structure in closed state of HSP90*α*-NTD was compared to the NTD extracted from full-length HSP90α (FL-HSP90-NTD) focusing on the ATP-lid segment that is stabilized in the closed conformation by the binding of ATP, co-chaperone and client protein^53^. The closed state of apo-HSP90α-NTD can be superimposed on the ATP-lid of dimeric full length HSP90α (PDB: 7L7J) with a RMSD of 4.1 Å (Cα backbone atoms). This RMSD value highlights the resemblance of the NMR-observed metastable excited state to the structure of the ATP-lid in FL-HSP90-NTD (Fig. 6c-d). In particular, the secondary structural motifs describing the ATP-lid segment are comparable: helix-3 and helix-4 are present, while the segment forming helix-5 in the open conformation is in an extended conformation in both FL-HSP90-NTD and the excited state of isolated HSP90-NTD. The presence of ATP in the structure of the full-length protein as well as the β-strand exchange leading to additional interactions may explain the remaining differences. Although our study was performed on isolated HSP90α-NTD, our results suggest that the functionally relevant closed state of HSP90 ATP-lid pre-exists in apo HSP90 before dimerization driven by ATP binding. Therefore, ATP binding selects and stabilizes a preexisting conformation before inducing further structural rearrangements. These results highlight the importance of investigating protein dynamics and structure at a physiologically relevant temperature. This is particularly relevant for structure-based design of new drugs against the ATP binding domain of human HSP90.

## Online Methods

### Preparation of isotopically labeled HSP90α-NTD samples

*E. coli* BL21-DE3-RIL cells transformed with a pET-28 plasmid encoding the N-Terminal domain of HSP90α from Homo Sapiens (HSP90α-NTD) with a His-Tag and a TEV cleavage site were progressively adapted in three stages over 24 h to M9/^2^H_2_O. In the final culture, bacteria were grown at 37°C in M9 medium with 99.85 % ^2^H_2_O (Eurisotop), 1 g/L ^15^N^1^H_4_Cl (Sigma Aldrich) and 2 g/L D-glucose-d_7_ or D-glucose-^13^C_6_-d_7_ for U-[^2^H, ^12^C, ^15^N] and U-[^2^H, ^13^C, ^15^N] HSP90α-NTD samples, respectively. For production of U-[^2^H, ^15^N, ^12^C], Ala-[^13^C^1^H_3_]^β^, Met-[^13^C^1^H_3_]^ε^, Leu-[^13^C^1^H_3_]^δ2^, Ile-[^13^C^1^H_3_]^δ1^, Thr-[^13^C^1^H_3_]^γ^, Val-[^13^C^1^H_3_]^γ2^ wild type HSP90α-NTD or mutants (R46A, R60A, D127A, S129A), HLAM-A^β^I^δ1^M^ε^(LV)^proS^T^γ^ labeling kits (NMR-Bio) were added to the M9/^2^H_2_O media^39^. To cross-validate sequence specific assignment of methyl probes, 56 single point mutant samples were produced^40^ (see supporting Fig. S2 for list of mutants).

For all samples, protein production was induced by 0.5 mM of IPTG when the O.D at 600 nm reached *ca.* 0.8, overnight at 20 °C. HSP90α-NTD samples were purified in two steps using a Ni-NTA affinity chromatography step followed by a size exclusion chromatography step as previously described^38^. For relaxation dispersion experiments, an additional tag cleavage step was performed for the preparation of the wild type HSP90α-NTD samples, by addition of TEV protease at room temperature for 2 h (in a ratio 1:10) before removal of His-tagged-TEV protease and cleaved His-Tag using reverse Ni-NTA affinity chromatography.

### NMR Spectroscopy

The final NMR buffer conditions were 20 mM HEPES (pH 7.5), 150 mM NaCl and 1 mM TCEP in either 100% ^2^H_2_O or 90% ^1^H_2_O/10% ^2^H_2_O. Single point mutant samples to cross-validate the assignment were concentrated at [0.1-0.4] mM. 40 µL of each sample was loaded in a 1.7 mm NMR tube and 2D ^1^H-^13^C SOFAST methyl TROSY^55^ experiments were recorded for an average duration of ∼ 1.5 h each, on a spectrometer equipped with a 1.7 mm cryogenically cooled, pulsed-field-gradient triple-resonance probe and operating at a ^1^H frequency of 850 MHz. All other NMR experiments were recorded on isotopically labeled HSP90α-NTD samples (200 µL of 0.5 mM labeled protein in 4 mm Shigemi tube) using a Bruker Avance III HD spectrometers equipped with 5 mm cryogenic probes.

3D HNH- and NNH-edited 3D NOESY experiments were recorded at 293 K on a spectrometer operating at a ^1^H frequency of 850 MHz using a U-[^2^H, ^13^C, ^15^N] wild type HSP90α-NTD sample in 90% ^1^H_2_O/10% ^2^H_2_O buffer. 3D CCH-edited HMQC-NOESY-HMQC NMR experiments^56,57^ were recorded on high field spectrometer, operating at a ^1^H frequency of either 850 MHz or 950 MHz, using U-[^2^H, ^15^N, ^12^C], Ala-[^13^C^1^H_3_]^β^, Ile-[^13^C^1^H_3_]^δ1^, Leu-[^13^C^1^H_3_]^δ2^, Met-[^13^C^1^H_3_]^ε^, Thr-[^13^C^1^H_3_]^γ^, Val-[^13^C^1^H_3_]^γ2^ samples of wild type, R46A-, R60A-, or S129A-HSP90α-NTD constructs in ^2^H_2_O buffer. Each 3D CCH-edited NOESY experiment was recorded over *ca.* 2-3 days, at 298 K for wild type HSP90α-NTD, 288 K for R46A- and S129A mutants or 293 K for R60A-HSP90α-NTD.

Methyl TROSY ^13^C^1^H-multiple quantum CPMG relaxation dispersion experiments^58^ modified with Ernst angle excitation^55^ experiments and ^15^N single quantum CPMG relaxation dispersion experiments were recorded at 293 K using U-[^2^H, ^15^N, ^12^C], Ala-[^13^C^1^H_3_]^β^, Ile-[^13^C^1^H_3_]^δ1^, Leu- [^13^C^1^H_3_]^δ2^, Met-[^13^C^1^H_3_]^ε^, Thr-[^13^C^1^H_3_]^γ^, Val-[^13^C^1^H_3_]^γ2^ WT- HSP90-NTD sample, on spectrometers operating at ^1^H frequencies of 700 MHz and 850 MHz (^13^C^1^H_3_ experiments) or of 600 MHz, 700 MHz and 850 MHz (^15^N experiments).. Methanol was used as a reference for precise temperature calibration on each spectrometer. The power of the ^13^C refocusing pulse in CPMG was set to 16 kHz (or 4 kHz for ^15^N), the CPMG relaxation period was set to 30 ms for ^13^C-(60 ms for ^15^N-) experiments. Dispersion profiles, recorded in an interleave manner, comprised 16 different CPMG frequencies (υ_cpmg_) and were ranging from 66 Hz to a maximum of 2000 Hz, whilst dispersion profiles comprised 11 different υ_cpmg_ and were ranging from 33 Hz to a maximum of 1000 Hz for ^15^N experiments. In addition, a reference spectrum was acquired for both types of experiments by omitting the relaxation period. Total experimental acquisition time for the relaxation dispersion experiment was 1 day for ^13^C-experiments and 2 days for ^15^N experiments.

### Structural Restraints

The following types of experimental distance restraints were used for structure calculations of HSP90-NTD variable segment from M98 to V136: (*i*) H_N_—H_N_ restraints derived from ^15^N-edited NOESY spectra acquired using a NOE mixing time of 240 ms; and (*ii*) methyl—methyl restraints from ^13^C-edited NOESY spectra. A qualitative approach was used in the derivation of distance restraints. In particular, intermethyl distance restraints were fixed between the barycenters of each proton triplets with distance bound of 2-4 Å and 4–6 Å for strong and weak cross-peaks detected in 3D-NOESY acquired with a short mixing time (*i.e.* 100 ms) or with distance bound of 4-8 Å for cross-peaks observed only in experiment acquired with longer NOE mixing time (*ca.* 300 to 400 ms). In the case of the mutant HSP90-NTD-R46A, intermethyl distance restraints were fixed between the barycenters of each proton triplets with distance bound of 2-8 Å as no short mixing 3D-NOESY experiment was acquired. H_N_—H_N_ distance bounds for observed NOE cross-peaks were fixed to 2–6 Å. For R60-D127 and R46-S129 stabilizing interaction identified using mutagenesis, supplementary distances restraints were fixed between oxygen acceptor atoms and arginine side chain nitrogen/hydrogen donor atoms with distance upper limits of 2/3 Å, respectively. Backbone chemical shifts of WT-HSP90-NTD constructs were used to determine φ and ψ dihedral angles using TALOS+ software^42,59,60^ and angular restraints with a tolerance of ±10° were applied. For the HSP90-NTD segment from M98 to V136, a total of 54 dihedral angles restraints were used for the calculation of WT- and R60A-HSP90-NTD structures. Due to widely broadened missing peaks, no angular restraints were experimentally derived from residues 105 to 115.

For the structure refinement of R46A-HSP90-NTD excited state, the upper limit of the 5 distance restraints specific to the ATP-lid closed state (Supporting Table S2) were reduced to 6.5 Å to take into account the lower population of this transiently sampled state, while the 22 structural restraints specific of the ATP-lid in open conformation (Supporting Table 2) were excluded of distance restraints set. The angular restraints for helix*−*5 (from I128 to V136) were excluded for the calculation R46A-HSP90-NTD structures, as we had evidences from initial structure calculation that helix*−*3 and helix-4 are preserved while helix*−*5 unfolds in the excited state (Supporting Table S5). Total number and distribution of structural restraints can be found in supporting Fig. S2 and Table S1.

### Structure calculation

Structural calculations were performed using CYANA 3.98.13 simulated annealing protocol^43^. The invariable segments from residues 11 to 97 and from 137 to 223 (Fig. 1) were treated as a rigid body core using a set of 13434 C_α_-C_α_ distance restraints and 477 dihedral angles (φ, ψ and χ_1_) extracted from X-ray crystal structure of apo HSP90-NTD (PDB code : 1YES^61^). The solution structures sampled by the variable segment of HSP90-NTD (Fig. 1) from residue M98 to V136 which harbors the specific loop covering the ATP-binding site was determined using NMR structural information obtained on 3 different samples (WT and R46A-, R60Amutants). Initial structure calculation of WT- and R46A-HSP90-NTD were performed using a two-state structure calculation protocol enabling each distance restraints to be satisfied in either ATP-lid open- or closed-state or both states simultaneously^54^.

Structural data acquired for the R60A-construct were used with a single state calculation protocol to refine the ATP-lid open structural ensemble (Fig. 3A). Conversely, the intermethyl NOE data set acquired for R46A-HSP90-NTD construct was used to refine the ATP-lid closed structural ensemble. For each HSP90-NTD structural state, a CYANA simulated annealing protocol using 25000 torsion angle dynamic steps^43^ was used to generate one thousand conformers starting from random coordinates and using NMR experimental restraints. The twenty conformers characterized by the lower target function values were selected for further refinement using a CNS restrained molecular dynamics protocol using a full Lenard-Jones potential and explicit water molecules^62^. In brief, the explicit solvent refinement consisted of the five following steps: (i) immersion in a 7.0 Å shell of water molecules and energy minimization; (ii) slow heating from 100 to 500 K in 100 K temperature steps with 200 MD steps per temperature step (time/step 3 fs), with harmonic position restraints on the protein heavy atoms that were slowly phased out during the heating stage; (iii) refinement at 500 K with 2,000 MD steps (time step 4 fs); (iv) slow cooling from 500 K to 25 K in 25 K temperature steps with 200 MD steps per temperature step (time step 4 fs); (v) final energy minimization (200 steps).

### Molecular Dynamics Simulations

The experimentally refined ensembles were used as starting points for MD simulations. Forty separate simulations were run for 1 µs, for the 20 different conformations observed for both the ATP-lid in the closed and open states.

#### 1.1 Setup and parameters

The all-atom Amber ff14SB force field^63^ was used with explicit water molecules described using TIP3P model^64^. The Gromacs software^65^ (single precision, 2020 versions) was used to perform preparation, equilibration and production steps. Each protein model was solvated in an octahedral box of about 13000 water molecules extending at least 13 Å away from the protein. Standard protonation states were assumed. The His189 was protonated on the Nδ1 atom, whereas the other three His residues were protonated on the Nε2 atom, in accordance with the pKa calculations performed by H++ server^66^ tested on 20 typical structures with relative permittivity of the solvent of 78, relative permittivity of the protein of 4 at a salt concentration of 0.17 mM. This resulted in a net charge of -11 for the protein. NaCl salt and Na+ counterions were added to obtain an electrically neutral system, with a salt concentration of about 170 mM. For each simulation, a steepest-descent energy minimization was used until the maximum force was lower than 100 kJ.mol^-1^.nm^-1^, followed by 100 ps constant-volume equilibration and 100 ps constant-pressure equilibration, both performed with heavy nonwater atoms restrained toward the starting structure with a force constant of 1000 kJ.mol^-1^.nm^-2^. Finally, 1020 ns were performed with constant pressure, constant temperature, without restraints. The first 20 ns were considered as an initial equilibration, and the last 1000 ns were used for the analysis presented in the article. All bonds involving hydrogen atoms were constrained to the equilibrium value using 1 iteration to the 4th order of the Linear Constraint Solver^67^, allowing for a time step of 2 ps for the leap-frog integrator. The temperature was kept constant at 300 K using two stochastic velocity rescaling thermostats^68^ for the solvent and for the protein, with a time constant of 2 ps. The pressure calculated with long-range correction was kept constant at 1 atm using a Parrinello-Rahman isotropic barostat^69^ with a relaxation time of 2 ps and a compressibility of 4.5×10^-5^ bar^-1^. Long-range electrostatics were handled by smooth particle-mesh Ewald (PME) summation^70^ with a fourth-order B spline interpolation and a grid spacing of 0.16 nm. The cutoff radius for Lennard−Jones interactions was initially set to 10 Å, but the neighbor list parameters were automatically optimized with an energy drift tolerance of 0.005 kJ.mol^-1^.ps^-1^ per particle^71^. The snapshots taken every 1 ns were analyzed with Gromacs and Mdanalysis^72^.

#### 1.2 Analysis of the Molecular Dynamics Simulations

##### 1.2.1 RMSD

For each of the 40 trajectories, 100 conformations were extracted (every 10 ns), yielding 4000 structures. The pairwise RMSD was calculated using the TTclust tool^73^. For each pair, the protein backbone atoms were superimposed on the rigid core of the protein (backbone of segments [11:97] and [137:223]), and the RMSD was calculated for the flexible ATP-lid, i.e., the backbone atoms for segment [98:136].

##### 1.2.2 Violations

Either for the closed or for the open state, the specific NOEs are defined by an ensemble of N couples of residues {ij} (see Table S2). At each time step, the distances d_ij_ between the methyles are measured, and the violation V relative to a given state is the average over the N violations v_{ij}_, i.e. V = 1/N ∑_{ij}_ v_{ij}_, where v_ij_ = max (0, d_ij_ -d_ij_^viol^). The distance d_ij_ between two protonated methyles is the distance between the two centers of masses of the three respective protons. We have chosen to use d_ij_^viol^ = <d_ij_> + 2 σ(d_ij_), where <d_ij_> and σ(d_ij_) are the average value and the standard deviation from the bundle of the 20 structures obtained after refinement under NMR restraints respectively. The tables S2A and S2B provides the details on the couples {ij}, the three protons defining the center of masses, and their respective d_ij_^viol^.

##### 1.2.3 Secondary structure

Analysis of the secondary structure per residue was done using DSSP. The analysis was done on 120 frames, extracted from the last 120 ns of the simulation^74^. The 3 helical types (α, 3_10_ and π) were considered as helices.

### Relaxation dispersion Data analysis

Peak intensities for relaxation-dispersion CPMG experiments were determined using the nonlinear fitting routine nlinLS from the NMRPipe software suite^75^. Relaxation dispersion profiles were generated from peak intensities, measured from pseudo 3D experiments using different CPMG frequencies and divided by intensities extracted from the reference spectrum^58^. After visual inspection to exclude too noisy relaxation dispersion curves, relaxation dispersion data of the non-overlapping peaks for which exchange contribution (R_ex_) to the transverse relaxation was ≥ 2 s^-1^ were analyzed and fitted with the software ChemEx^76^ using a two-state exchange model. Errors for R_2,eff_ rate values were estimated from twice the noise measured in the spectra. However, when errors were less than 2% of the R_2,eff_ rate value, an error of 2% was assumed^58^.

First, a grid search using relaxation data was carried out where both conformational exchange rate and population of the excited state were fixed (Supporting Fig S7) to identify the minimum in order to select these parameters as starting parameters for the global numerical fitting (a single and identical local minimum was identified for both ^15^N and ^13^CH_3_ relaxation datasets). For each experimental temperature, a global numerical fit of ^13^CH_3_-CPMG relaxation dispersion profiles was performed to obtain global exchange parameters (exchange rate (k_ex_) and the population of the excited state (P*)) as well as residue specific parameters (absolute chemical shift difference between the ground and the excited state of the system under chemical exchange, |Δω_C_| and |Δω_H_|). A jackknife resampling was run to ensure that all residues globally fitted were part of the same exchange process and finally 40 Monte-Carlo repeats were used to estimate precision on global fit results. Experimental data for 21 different methyl probes were used for the global fitting of the methyl ^13^C-^1^H CPMG-RD experiments acquired at 293 K.

### Data Availabilities

All the experimental NMR data used for the structural and dynamics characterization of human HSP90-NTD as well as structures ensembles together with corresponding structural restraints used for the calculation of HSP90-NTD in ATP-Lip open (R60A mutant) and closed (R46A mutant) will be deposited in BMRB and PDB databases and accession codes will be provided before publication.

## Acknowledgements

The authors thank Drs. R. Awad and C. Laguri for help and advices. This work used the high field NMR and isotopic labeling facilities at the Grenoble Instruct-ERIC Center (ISBG; UAR 3518 CNRS-CEA-UGA-EMBL) within the Grenoble Partnership for Structural Biology (PSB). Platform access was supported by FRISBI (ANR-10-INBS-05-02) and GRAL, a project of the University Grenoble Alpes graduate school (Ecoles Universitaires de Recherche) CBH-EUR-GS (ANR-17-EURE-0003). IBS acknowledges integration into the Interdisciplinary Research Institute of Grenoble (IRIG, CEA). This work was supported by grants from CEA/NMR-Bio (research program C24990), by the *Region Auvergne-Rhône-Alpes* (pack ambition recherche 2019 - P089), by the French National Research Agency in the framework of the “*Investissements d’avenir*” program (ANR-15-IDEX-02) and by a fellowship (to F.H.) from “La Ligue contre le Cancer”.

## Author contributions

C.L., E.R., F.H., J.B., M.F. designed the experiments; E.C. and F.H. prepared the samples; F.H., J.B., B.B. and P.M. set up and collected the NMR experiments; E.R., F.H., J.B., and P.G. analyzed the NOESY experiments; A.F. and E.R. performed structural calculations; F.H., J.B. and P.M. analyzed the CPMG experiments; C.L., E.R., and P.J. performed and analyzed the MD simulations; C.L., E.R., F.H. and J.B. wrote the manuscript. All authors discussed the results, corrected the manuscripts and approved the final version.

