## Supplementary Information for "Visualizing the Transiently Populated Closed-State of Human HSP90 ATP Binding Domain"

### Supporting Information

Faustine Henot[1,#], Elisa Rioual[1,2,#], Adrien Favier[1], Pavel Macek[1,3], Elodie Crublet[3], Pierre Josso[2], Bernhard Brutscher[1], Matthias Frech[4], Pierre Gans[1], Claire Loison[2,*], Jerome Boisbouvier[1,*]

**Affiliation:**
1. Univ. Grenoble Alpes, CNRS, CEA, Institut de Biologie Structurale (IBS), 71, avenue des martyrs, F-38044 Grenoble, France.
2. Institut Lumière Matière, University of Lyon, Université Claude Bernard Lyon 1, CNRS, F-69622, Villeurbanne, France.
3. NMR-Bio, 5 place Robert Schuman, F-38025 Grenoble, France.
4. Discovery Technologies, Merck KGaA, Frankfurter Straße 250, 64293 Darmstadt, Germany.
### These authors contributed equally to this work

#### Table of Contents

Table S1: Summary of structural restraints. p. 2

Table S2: Definition of NOE-derived distance restraints specific of a single state. p. 3

Table S3: Summary of NMR refined structure statistics. p. 4

Table S4: Identification of characteristic distances of dimeric forms of HSP90α-NTD. p. 5

Table S5: NMR-based structural calculations supporting the absence of helix-5 in ATP-lid closed state. p. 6

Figure S1: Assigned 2D NMR spectra of HSP90α-NTD. p. 7

Figure S2: HSP90α-NTD assignment. p. 8

Figure S3: Summary of NOEs distance restraints. p. 9

Figure S4: Assigned 2D $^{1}H$-$^{13}C$ SOFAST methyl TROSY of HSP90α-NTD mutants p. 10

Figure S5: Intensity variation of characteristic NOEs. p. 11

Figure S6. Time evolution of the violations of the NOE-derived distance restraints specific of either the open or the closed conformation for the 20 non-restrained molecular dynamics simulations starting from ATP-lid closed state conformers. p. 12

Figure S7: Analysis of HSP90-NTD dynamics using relaxation dispersion experiments. p. 14

Figure S8: Computed Evolutionary Couplings (ECs) for HSP90-NTD. p. 15

Figure S9: Probability of helical structures along the sequence of the ATP lid, in 4 MD simulations depicting a transition of the ATP-lid from a closed to an open state. p. 16

**Table S1: Summary of structural restraints used for structure calculation of WT-, R60A- and R46A-HSP90α-NTD.**

| | | WT[(a)] | R60A | R46A |
|---|---|---|---|---|
| **Experimental distance restraints** | | | | |
| **$CH_3$-$CH_3$** | | **111** | **114** | **81[(b)]** |
| Sequential / local (≤i+4) / long range / ambiguous | | 4 / 12 / 75 /20 | 5 / 13 / 78 / 18 | 4 / 15 / 57 /5 |
| **$H_N$-$H_N$** | | **41** | **41** | **41** |
| Sequential / local (≤i+4) / long range / ambiguous | | 18 / 19 / 3 /1 | 18 / 19 / 3 / 1 | 18 / 19 / 3 / 1 |
| **H-bonds** | | **0** | **2** | **2** |
| **Experimental angular restraints** | | | | |
| **Dihedrals** | φ, ψ | **36+54[(a)]** | **54** | **36[(c)]** |
| **Restraints for rigid body** | | | | |
| Dihedrals | φ, ψ | 475 + 477[(a)] | 477 | 475 |
| Distances | $C_\alpha$-$C_\alpha$ | 13434 + 13434[(a)] | 13434 | 13434 |

**(a)** Structure calculations using NMR restraints derived from WT-HSP90-NTD sample were performed using a two-state structure calculation protocol enabling each experimental distance restraint to be satisfied in either ATP-lid open- or closed-state or both states simultaneously[54]. The angular restraints, and distance restraints for the rigid body core (segments 11-97 and 137-223) were applied on both states simultaneously. Dihedral restraints, corresponding to helix-5 in the ATP-lid open state (from residues 128 to 136), were not applied for the closed ATP-lid state (See note (c) for further information).
**(b)** A total of 103 inter-methyl distance restraints were experimentally extracted using R46A-HSP90α-NTD sample to characterize the ATP-lid segment (from residue M98 to V136). Removal of R46/S129 hydrogen bond by mutagenesis leads to an increase of the population of the closed state, but NOEs corresponding to an ATP-lid in both open and closed conformations were detected simultaneously. Therefore, we used the previously refined solution structure with the ATP-lid in an open position (obtained using R60A-HSP90α-NTD sample) to identify and exclude 22 NOEs specifically assigned to the ATP-lid in an open state (see supporting Table S2). The remaining 81 NOEs were completed with backbone restraints previously obtained on WT-HSP90α-NTD sample for the final structure calculation using a single state structure calculation protocol.
**(c)** Structure calculation allowed us to identify large violations due to incompatibilities between helix-5 dihedral restraints and NOEs specific of the ATP-lid closed state (Supporting Table S5). Therefore, the angular restraints from residues 128 to 136 were omitted for the final CYANA structure calculation of ATP-lid closed state using restraints derived from the R46A-HSP90α-NTD sample.

#### Table S2: Definition of NOE-derived distance restraints specific of a single state.

**A)** NOE-derived distance restraints specific of the ATP-lid closed state. NOEs observed only for the WT- and R46A-HSP90α-NTD mutants and not for R60A-HSP90α-NTD sample. In the structure ensembles calculated with a two-state protocol using distance restraints derived from NOEs detected on WT- and R46A-HSP90-NTD, those restraints are satisfied only in the ATP-lid closed state and violated by more than 5 Å in the ATP-lid open state.

| Restraint | Residue i | Pseudoatom i | Residue j | Pseudoatom j | $d_{ij}^{max}$ (Å)[a] | $d_{ij}^{viol}$ (Å)[b] |
|---|---|---|---|---|---|---|
| **1** | LEU64 | HD2* | MET130 | HE* | 6.5 | 10.08 |
| **2** | THR65 | HG2* | ALA121 | HB* | 6.5 | 10.43 |
| **3** | THR65 | HG2* | ALA124 | HB* | 6.5 | 9.85 |
| **4** | THR65 | HG2* | ALA126 | HB* | 6.5 | 10.19 |
| **5** | THR65 | HG2* | MET130 | HE* | 6.5 | 10.19 |

**B)** NOE-derived distance restraints specific of the ATP-lid open state. NOEs observed for WT-, R46A-, or R60A-HSP90α-NTD samples. In the structure ensembles calculated with a two-state protocol using distance restraints derived from NOEs detected on WT- and R46A-HSP90-NTD, those distance restraints are satisfied only in the ATP-lid open state and violated by more than 5 Å in the ATP-lid closed state.

| Restraint | Residue i | Pseudoatom Pi | Residue j | Pseudoatom Pj | $d_{ij}^{max}$ (Å)[a] | $d_{ij}^{viol}$ (Å)[b] |
|---|---|---|---|---|---|---|
| 1 | ALA24 | HB* | MET119 | HE* | 8 | 11.87 |
| 2 | ILE26 | HD1* | THR115 | HG2* | 8 | 5.54 |
| 3 | ILE26 | HD1* | MET119 | HE* | 8 | 8.59 |
| 4 | ILE26[c] | HD1* | ILE131[c] | HD1* | 8 | [f] |
| 5 | LEU29 | HD2* | THR115 | HG2* | 8 | 6.63 |
| 6 | LEU29 | HD2* | MET119 | HE* | 8 | 6.16 |
| 7 | LEU29 | HD2* | LEU122 | HD2* | 8 | 8.89 |
| 8 | LEU32 | HD2* | MET119 | HE* | 8 | 4.52 |
| 9 | LEU32 | HD2* | LEU122 | HD2* | 8 | 6.24 |
| 10 | LEU32[c] | HD2* | MET130[c] | HE* | 8 | [f] |
| 11 | LEU32 | HD2* | ILE131 | HD1* | 8 | 9.65 |
| 12 | THR36 | HG2* | LEU122 | HD2* | 8 | 8.53 |
| 13 | THR36 | HG2* | ILE131 | HD1* | 8 | 11.65 |
| 14 | ILE43 | HD1* | LEU122 | HD2* | 8 | 9.42 |
| 15 | ILE43[d] | HD1* | MET130[d] | HE* | 8 | [f] |
| 16 | ALA111 | HB* | MET119 | HE* | 8 | 10.55 |
| 17 | THR115 | HG2* | VAL136 | HG2* | 8 | 6.64 |
| 18 | MET119 | HE* | VAL136 | HG2* | 8 | 8.76 |
| 19 | LEU122 | HD2* | VAL136 | HG2* | 8 | 8.31 |
| 20 | LEU122 | HD2* | LEU143 | HD2* | 8 | 7.98 |
| 21 | LEU32 | HD2* | THR115 | HG2* | 8 | 10.72 |
| 22 | LEU32 | HD2* | ALA126 | HB* | 8 | 8.53 |
| 23 | MET119[e] | HE* | ILE131[e] | HD1* | 8 | [f] |
| 24 | LEU122[e] | HD2* | ILE131[e] | HD1* | 8 | [f] |

[a] For each NOE cross-peak observed associated to a pair of residue {i,j}, $d_{ij}^{max}$ defines the upper limit for the distance restraints applied for structure calculations.
[b] In the MD simulations, the violation relative to a given NOE-derived distance restraint differs from zero if the distance between the pseudoatoms Pi and Pj is larger than $dij^{viol}$ (see more details Section 1.2.2 of Online Methods).
[c] NOEs not observed in the construct HSP90-R46A
[d] NOEs not observed in the construct HSP90-R60A
[e] NOEs not observed in the construct HSP90-WT
[f] Distance not used to calculate violation in the analysis of the MD trajectories.

**Table S3: Summary of NMR refined structure statistics for HSP90-NTD in ATP-lid open and closed states.**

| | Closed state (R46A-HSP90-NTD) | Open state (R60A-HSP90-NTD) |
|---|---|---|
| **RMSD to mean structure**[a] | | |
| *Backbone atoms (Å)* | 1.52 ± 0.28 | 0.86 ± 0.18 |
| *Heavy atoms (Å)* | 2.08 ± 0.29 | 1.30 ± 0.21 |
| **Distance restraints violations** | | |
| *Average number of violations (>0.4Å)* | 0 | 0 |
| *Highest violation (Å)* | 0.37 | 0.24 |
| **Ramachadran analysis**[b] | | |
| *Most favored (%)* | 87.8 | 90.0 |
| *Additionally allowed region (%)* | 10.7 | 9.0 |
| *Generously allowed region (%)* | 1.3 | 0.9 |
| *Disallowed region (%)* | 0.2 | 0.1 |

[a]RMSD to the mean structure for the ATP-lid (from the residue 98 to 136) calculated for the 20 structures with the lower CYANA target functions refined using CNS restrained molecular dynamics.
[b]Ramachadran statistics calculated with PROCHECK for both states of HSP90-NTD.

#### Table S4: Identification of characteristic distances of dimeric forms of HSP90α-NTD.

Close by pairs of methyl groups representative of a possible dimerization of HSP90-NTD with the ATP-lid closed state and intermolecular exchange of the two $\beta_1$ strands are shown column 1. Such pair of methyl groups correspond to inter-chain putative NOEs involving one methyl group from residue 13 to 41 and any other methyl group of the NTD for which the measured distance was ≤ 11 Å in the full-length dimer HSP90α in closed form (PDB: 7L7J) but > 11 Å for both the solution structure ensemble representative of the ground/open and excited/closed ATP-lid states (Fig. 3). In the second column, presence/absence of putative specific NOEs of HSP90α-NTD dimer have been investigated. Their presence/absence has been verified in the 3D $^{13}CH_3$-edited NOESY spectrum acquired using the methyl-labeled R46A-HSP90α-NTD sample.

| **Contacts ($d_{HH}$ ≤ 11 Å) specific of dimeric full-length HSP90α** | **Present in NOESY spectra of R46A-HSP90-NTD ?** |
|---|---|
| 17 - 149 | No |
| 17 - 174 | No |
| 17 - 176 | No |
| 21 - 26 | No |
| 21 - 110 | No |
| 24 - 29 | No |
| 24 - 30 | No |
| 24 - 33 | No |
| 24 - 110 | No |
| 24 - 136 | No |
| 29 - 109 | No |
| 32 - 111 | No |
| 33 - 24 | No |
| 33 - 111 | No |
| 36 - 111 | No |
| 36 - 136 | No |

#### Table S5: NMR-based structural calculations supporting the absence of helix-5 in ATP-lid closed state.

All CYANA structural calculations performed without angular restraints for helix-5, but including all experimental NOE-derived distance restraints have a target function below 5 $kJ.mol^{-1}$ and no violation of NMR experimental restraints. In contrast, when dihedral restraints are introduced to force the formation of helix-5, the target function increases above 6 $kJ.mol^{-1}$ and 3 to 4 violations of NMR structural restraints are observed.

| Helices present in the structural calculation | Energy ($kJ.mol^{-1}$) | Distance restraint violation | Angular restraint violation |
|---|---|---|---|
| None | 4.51 ± 0.04 | ⌀ | ⌀ |
| α3 | 4.54 ± 0.05 | ⌀ | ⌀ |
| α4 | 4.60 ± 0.04 | ⌀ | ⌀ |
| α5 | 6.13 ± 0.06 | 2[a] | 1[b] |
| α3α4 | 4.56 ± 0.02 | ⌀ | ⌀ |
| α3α5 | 6.13 ± 0.08 | 2[a] | 1[b] |
| α4α5 | 6.23 ± 0.09 | 2[a] | 2[c] |
| α3α4α5 | 6.14 ± 0.07 | 2[a] | 2[c] |

[a] Violated distance restraints : THR65 HG2* - MET130 HE* and ILE131 HD1* - LEU143 HD2*
[b] Violated angular restraints: PHI SER211
[c] Violated angular restraints : PHI SER211 and CHI1 ILE214
⌀ : no violation

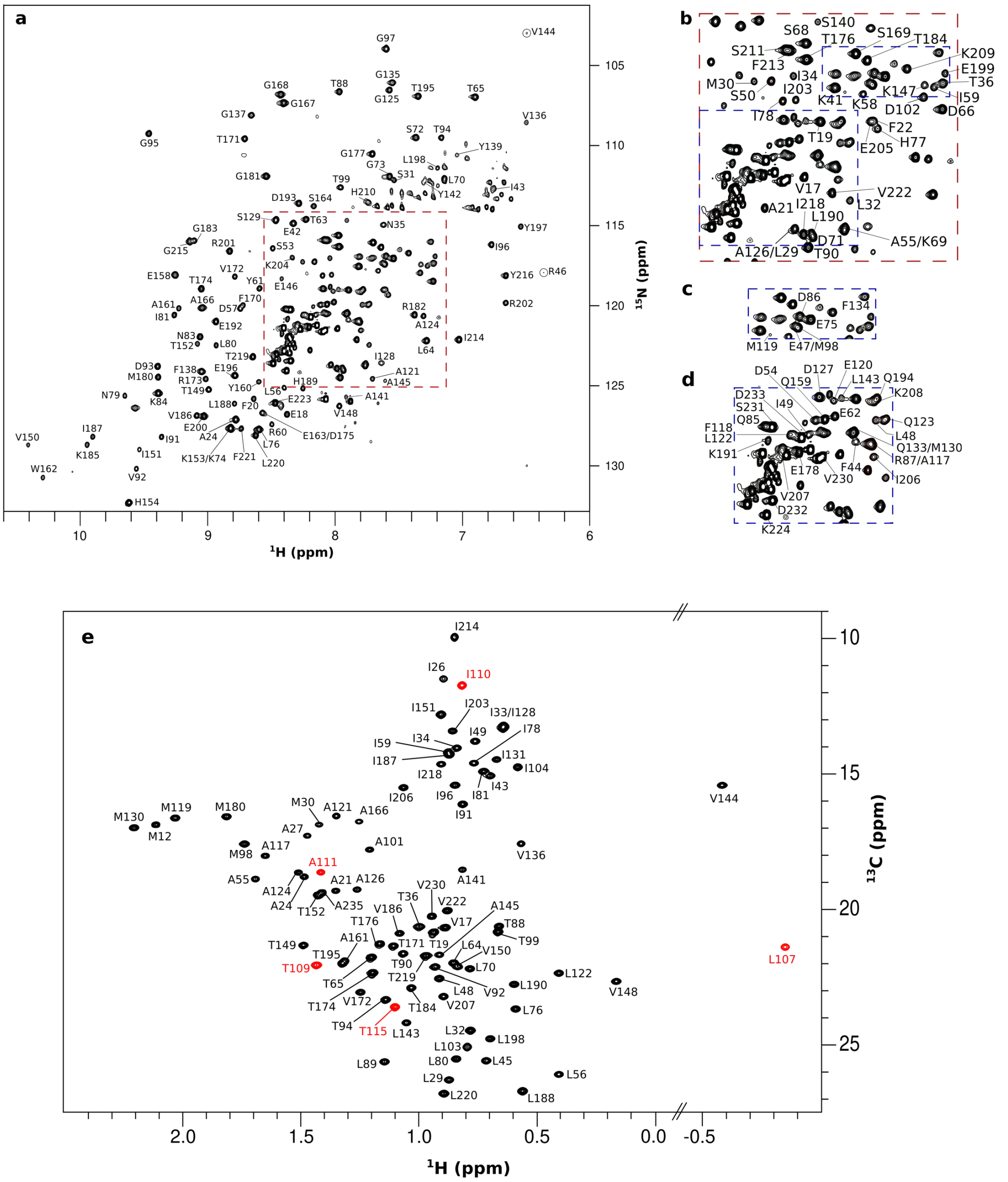


**Figure S1: Assigned 2D NMR spectra of HSP90α-NTD**. **a)** $^{15}$N-BEST-TROSY spectrum of HSP90α-NTD in its apo state, acquired at 293 K and at a $^{1}$H frequency of 600 MHz. Panels **b-d)** present zooms of 2D $^{15}$N-BEST-TROSY spectrum. Each assigned signal (a, b, c, d) is annotated with the corresponding residue number[36-38]. Panel **e)** displays the fully assigned 2D $^{1}$H-$^{13}$C SOFAST methyl TROSY spectrum of HSP90α-NTD apo, recorded at 298 K. Each signal is annotated with the corresponding residue number[38]. In red are displayed residues belonging to the stretch [105-116] invisible on $^{15}$N-BEST-TROSY due to conformational exchange in the µs-ms timescale.

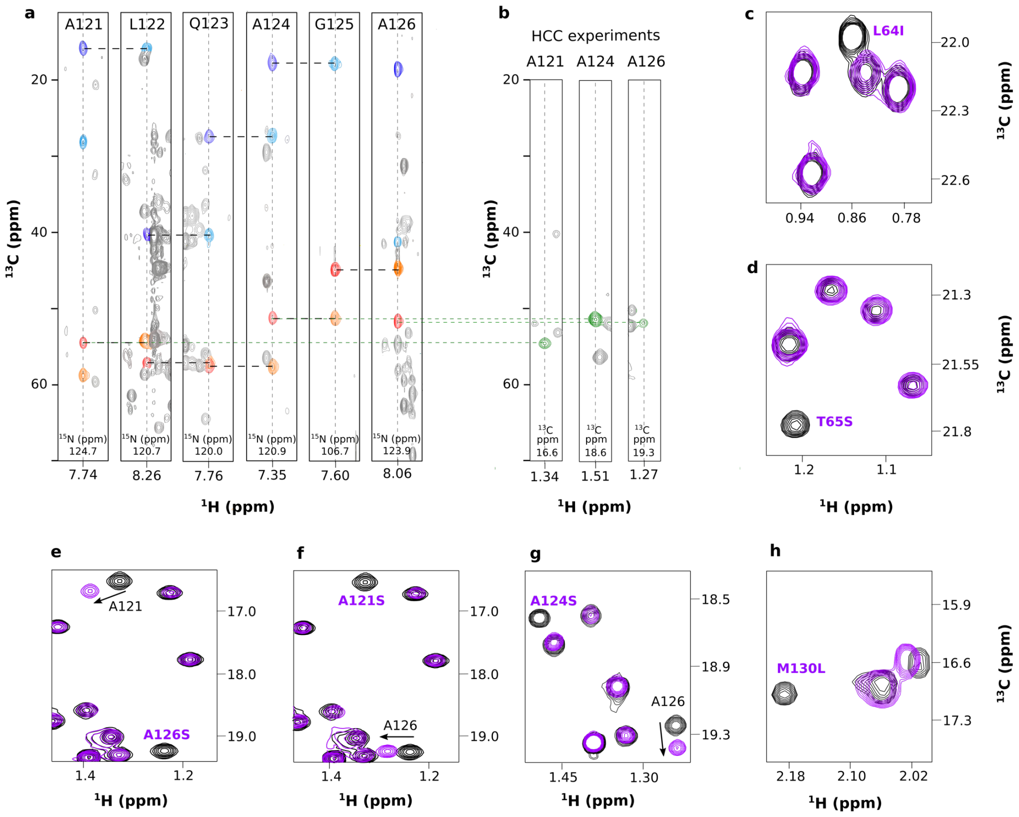


**Figure S2: HSP90α-NTD assignment. a)** Examples of 2D-extracts from 3D HNCA, HN(CO)CA, HNCACB, HN(CO)CACB showing backbone correlations from A121 to A126. Spectra were acquired at 298 K on a NMR spectrometer operating at a $^{1}$H frequency of 600 MHz using a U-[$^{2}$H, $^{13}$C, $^{15}$N]-HSP9α-NTD sample. $C\alpha_{(i)}$, $C\alpha_{(i-1)}$, $C\beta_{(i)}$, $C\beta_{(i-1)}$, are colored in red, orange, dark blue and light blue, respectively. **b)** Examples of 2D-extracts from 'out and back' HCC experiment to transfer assignment from backbone atoms to the methyl groups**. c-h)** Cross-validation of assignment of methyl groups using mutagenesis. Examples of 2D $^{1}$H-$^{13}$C SOFAST methyl TROSY spectra of 6 HSP90α-NTD mutants. Each mutant spectrum extract (purple) is superimposed with the wild type protein (black). A total of 56 single point amino acid mutations were generated by GeneCust and for each of these samples a single type of methyl group was labeled by addition of the corresponding NMR-Bio kit (SLAM-$A^{\beta}$, SLAM-$M^{\varepsilon}$, SLAM-$I^{\delta 1}$, SLAM-$T^{\gamma}$, SLAM-$V^{proS}$ or DLAM-$LV^{proS}$) in M9/$^{2}H_2O$ media. The following mutants were produced and analyzed by NMR: M12L, M30L M119L, M130L, M180L, A21S, A24S, A27S, A55S, A101S, A111S, A117S, A121S, A124S, A126S, A161S, A166S, A235S, T19S, T36S, T65S, T99S, T109S, T115S, T152S, T171S, T174S, T184S, T195S, T219S, I26V, I33V, I34V, I81V, I104V, I110V, I128V, I131V, I214V, V17I, V136I, V150I, V172I, V186I, V222I, V230I, L29I, L32I, L45I, L48I, L56I, L64I, L70I, L107A, L122I, L143I).

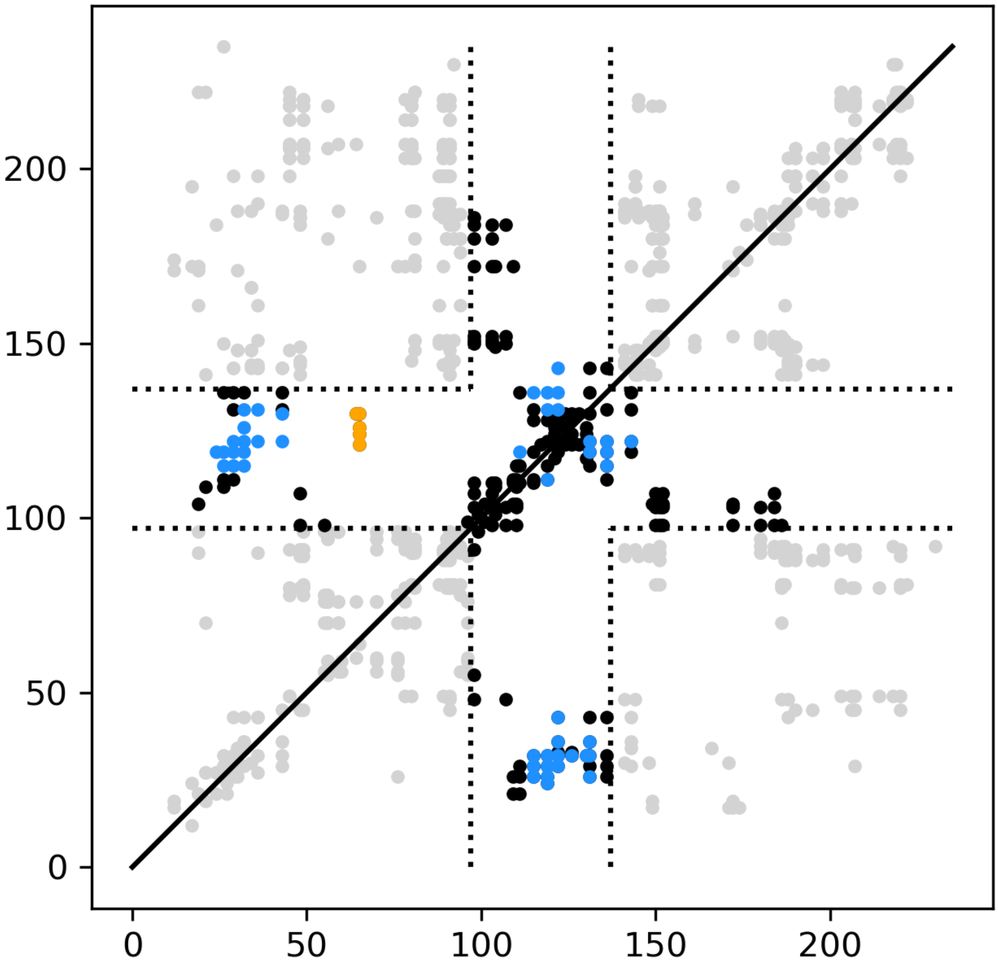


**Figure S3: Summary of NOEs distance restraints. a)** Experimentally detected $CH_3$-$CH_3$ (top left) and $H_N$-$H_N$ (bottom right) NOEs in WT-HSP90α-NTD. **b)** Experimentally detected $CH_3$-$CH_3$ NOEs for R46A- (top left) and R60A- (bottom right) WT-HSP90α-NTD. Black points represent NOEs of the ATP-lid present in both states. Blue/orange points represent NOEs specific of ATP-lid in open/closed states, respectively (Supporting Table S2).

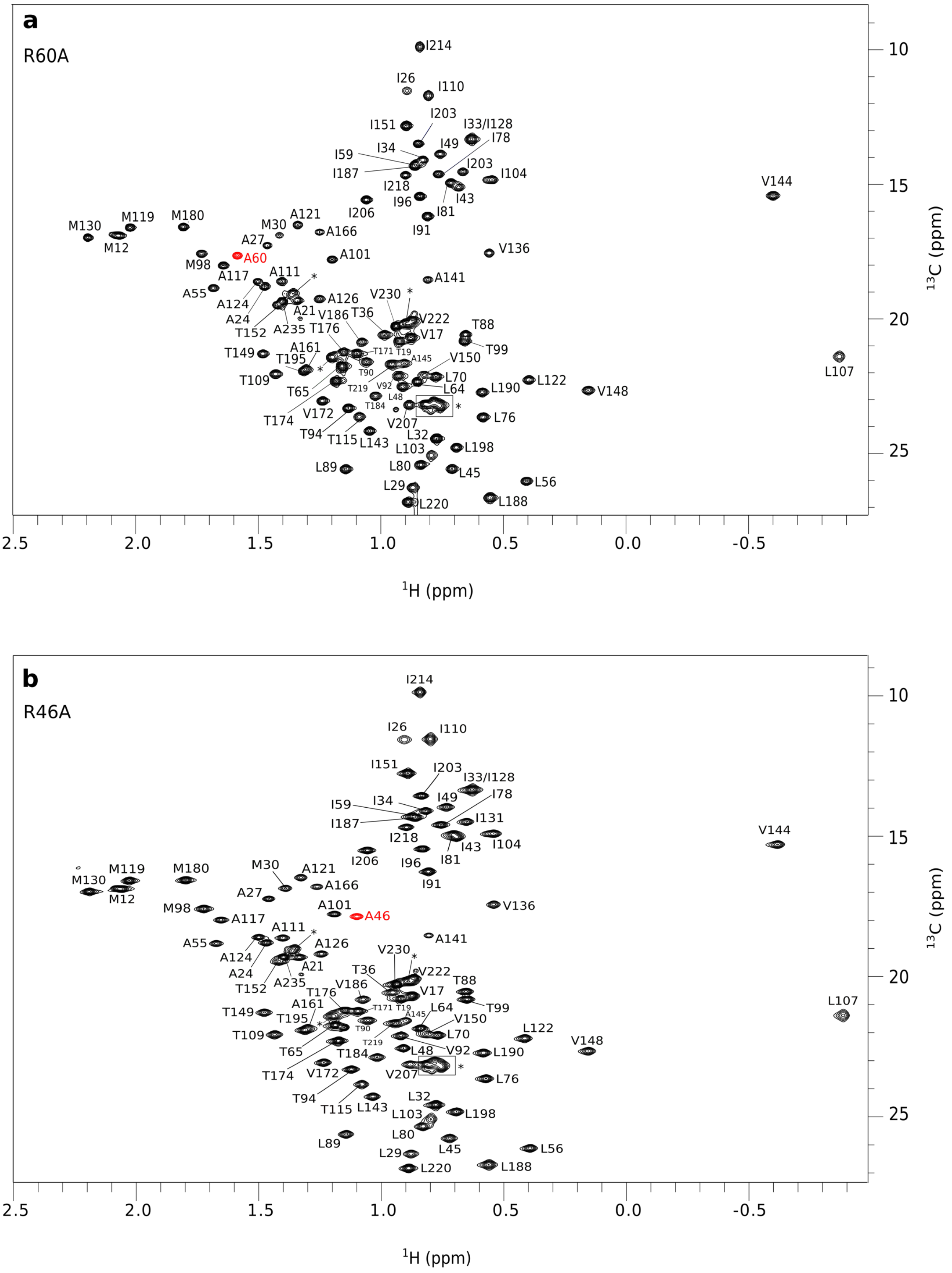


**Figure S4: Assigned 2D $^{1}$H-$^{13}$C SOFAST methyl TROSY of HSP90α-NTD mutants. a)** R60A- and **(b)** R46A-HSP90α-NTD mutants, recorded at 293 K. Each signal is annotated with the corresponding residue number. Peaks annotated with the symbol * are peaks belonging to the uncleaved N-terminal tag. Extra alanine methyl is colored in red.

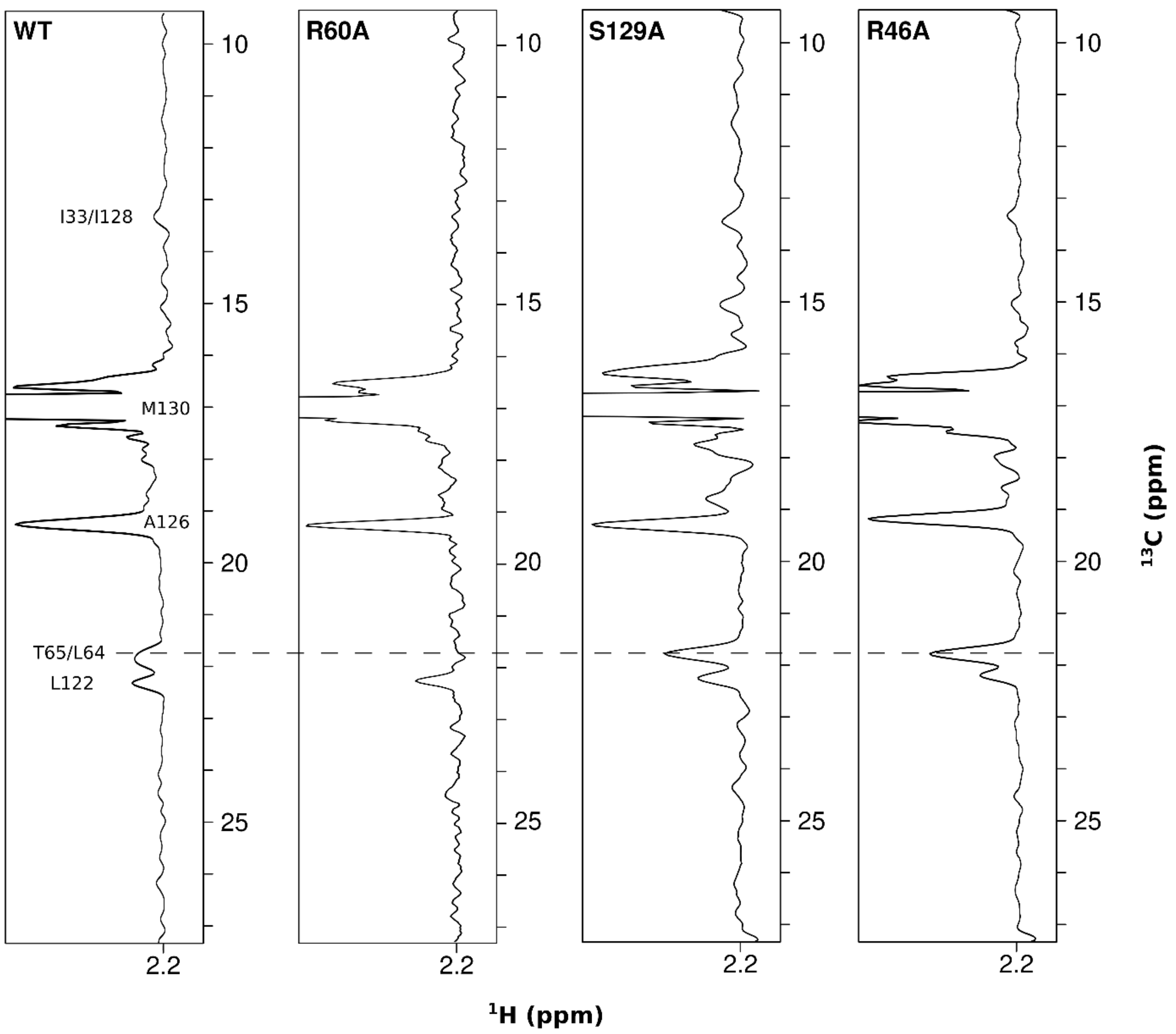


**Figure S5: Intensity variation of characteristic NOEs.** 1D traces extracted from 3D CCH HMQC-NOESY-HMQC experiments showing M130 as diagonal peak and associated NOEs cross peaks for the wild type protein HSP90α-NTD and three mutants: R60A, S129A and R46A. 3D $^{13}$C-edited NOESY spectra were acquired during *ca.* 2 days each using high-field NMR spectrometers operating at 950 or 850 MHz and at a temperature of 25°C (WT), 20°C (R60A) or 15°C (R46A), taking into account the lower stability of mutants. NOE mixing time were optimized to delays ranging from 0.3 (15°C) to 0.4 s (25°C), corresponding to the estimated maximum intensities of NOEs buildups for each experimental condition.

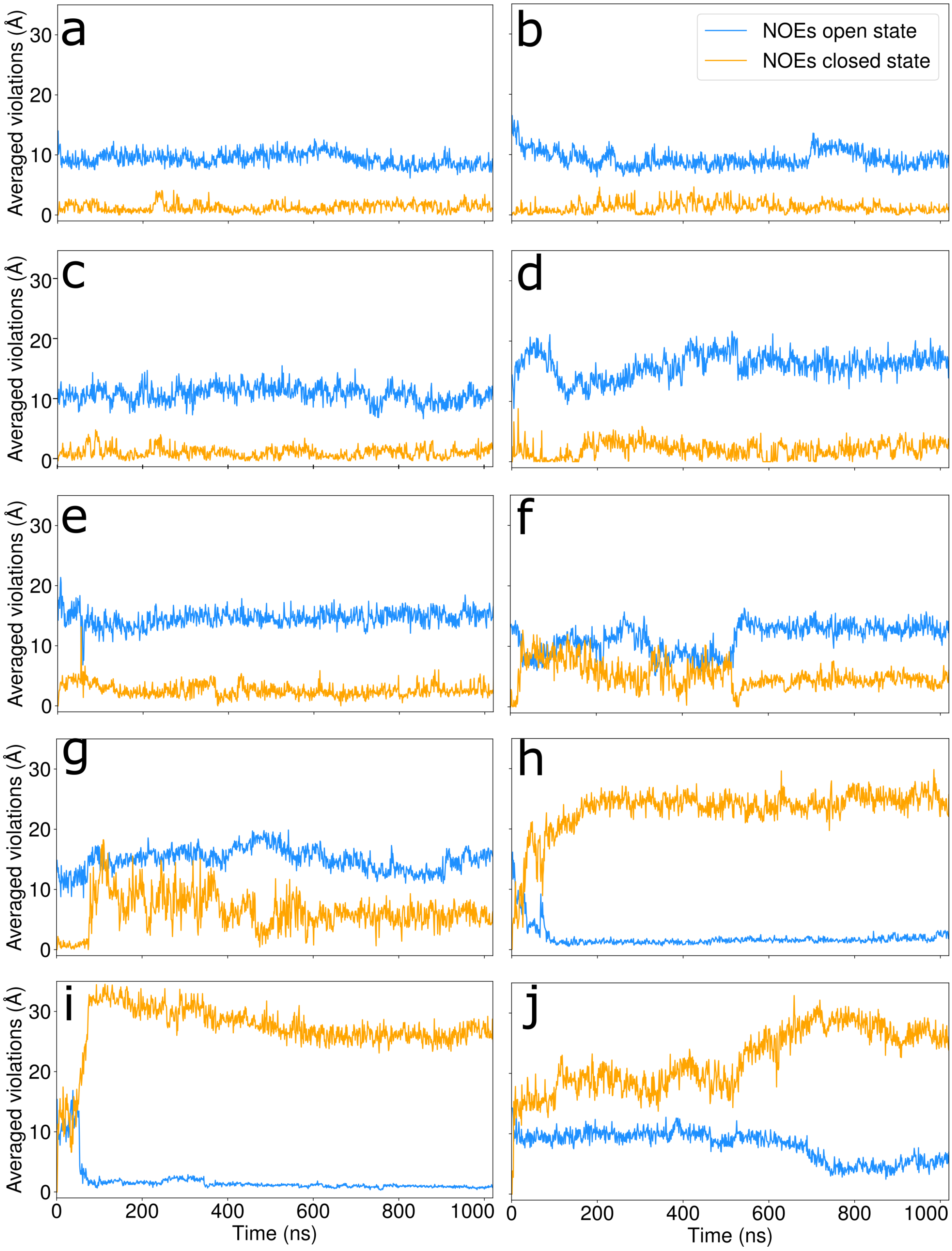


**Figure S6a. Time evolution of the violations of the NOE-derived distance restraints specific of either the open or the closed conformation (orange and blue respectively), for the 20 non-restrained molecular dynamics simulations starting from ATP-lid closed state conformers.** The violations are defined in section 1.2.2 of Online Methods, and the simulations were classified according to the violations. **a-c**: relatively stable trajectories, compatible with closed-state-NOE distances during several hundreds of nanoseconds. **d-g**: relatively stable trajectories, remaining nearby the closed state during several hundreds of nanoseconds. **h-j**: unstable trajectories with transition toward a conformation close to the open-state.

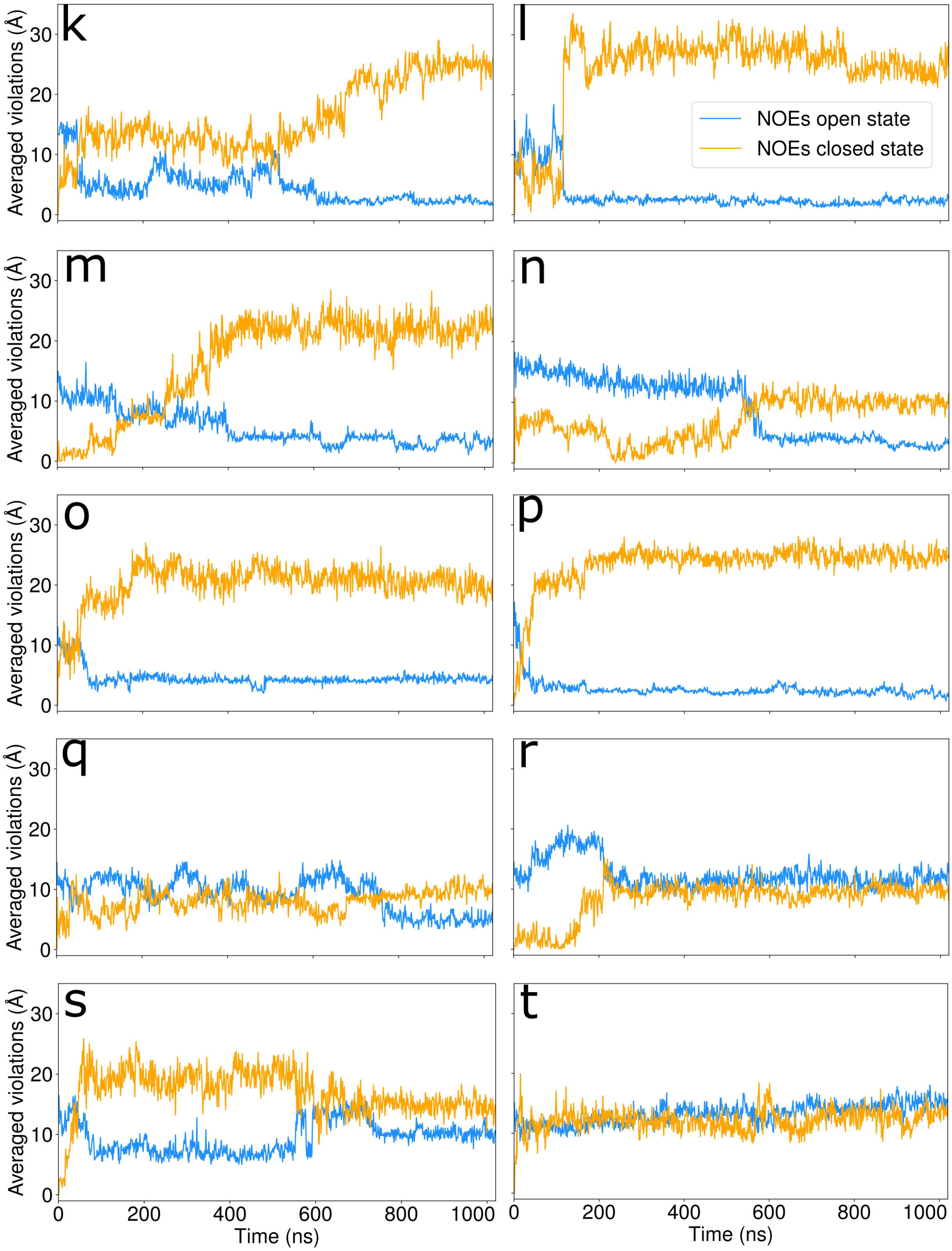


**Figure S6b : same as Figure S6a. k-p**: unstable trajectories with transition toward a conformation close to the open-state. **q-t**: unstable trajectories with transition toward a conformation neither closes nor open-state.

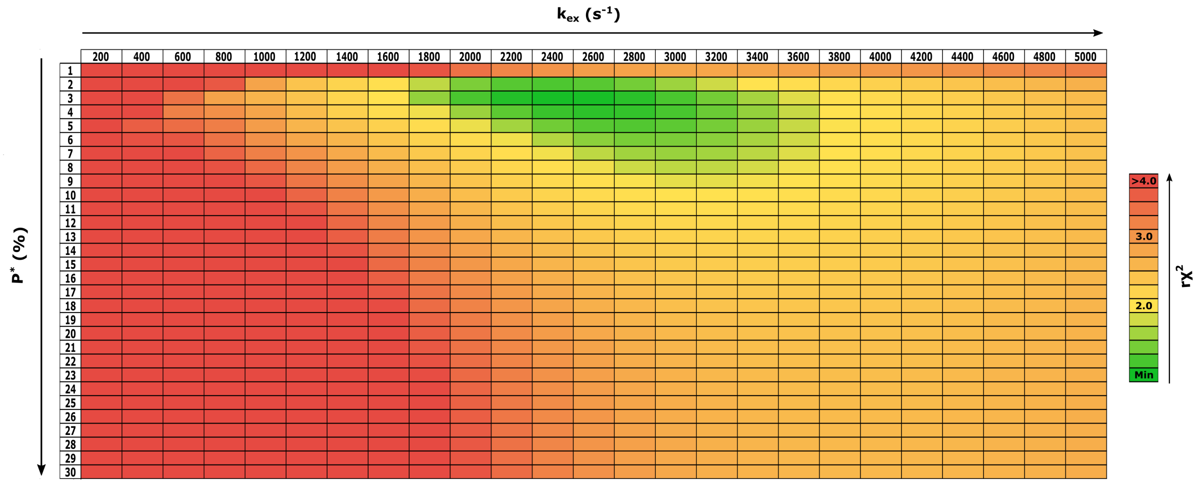


**Figure S7: Analysis of HSP90-NTD dynamics using relaxation dispersion experiments.** Grid search table displaying the reduced $\chi^2$ of the fit obtained from $^{13}$C-relaxation data acquired at 293 K, when exchange rate and population of the excited state were fixed. Going from green to red the reduced $\chi^2$ values increase.

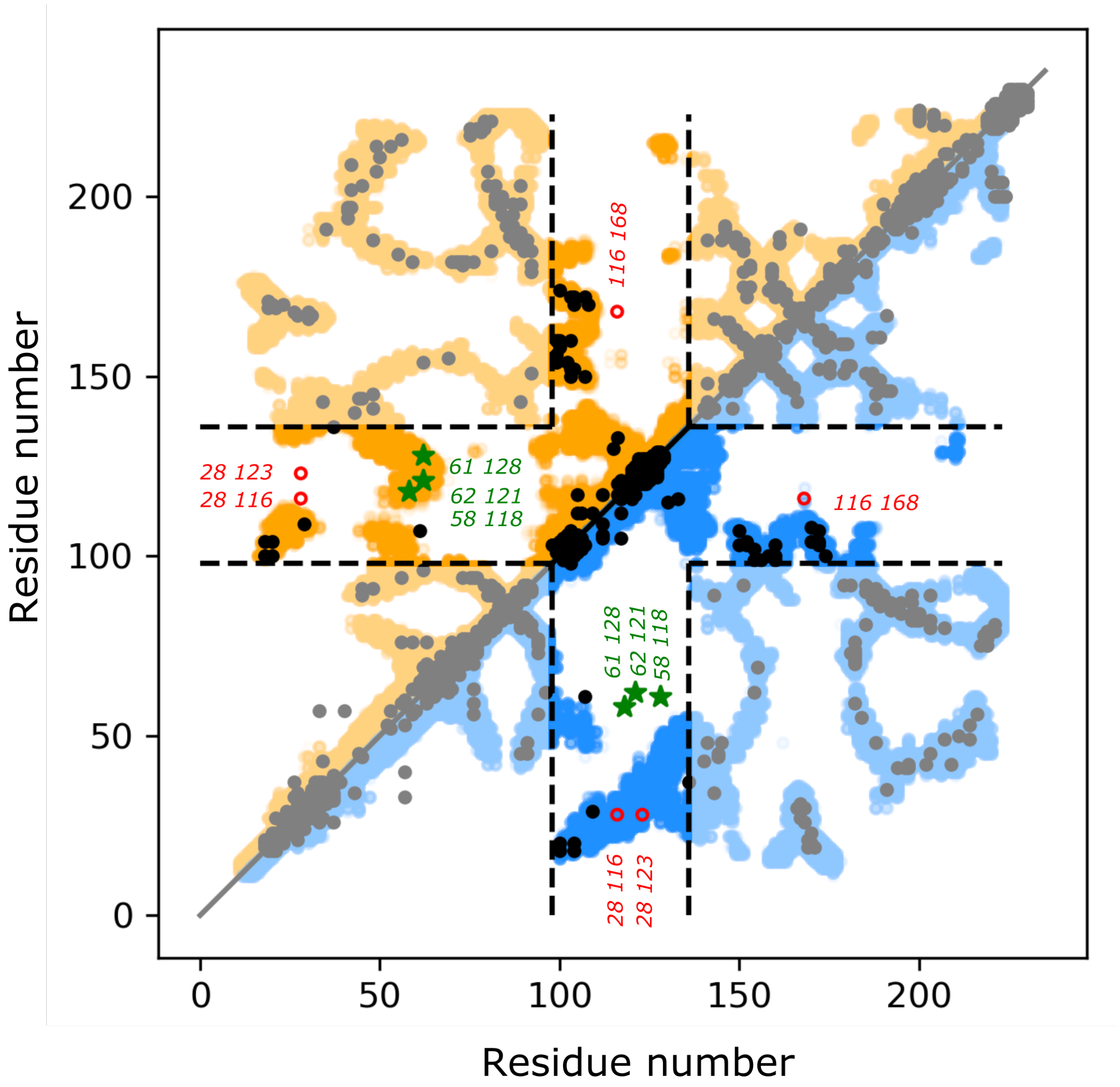


**Figure S8: Computed Evolutionary Couplings (ECs) for HSP90-NTD.** Contact map of top-ranked predicted ECs (black dots) overlaid on HSP90-NTD contact maps of ground/open (blue) and excited/closed (orange) ATP-lid states. 453 highly probable ECs ($p > 0.85$) predicted by EVcouplings (https://v2.evcouplings.org) were computed[47] starting from an alignment of 4414 homologue sequences of HSP90-NTD coming from the data base UniRef90. 71 ECs involving at least one residue of the HSP90 protein segment [98-136] are highlighted in the middle of the matrix. The contact maps have been plotted using both ground/open and excited/closed structures ensembles (Fig. 3). A colored dot was plotted for each $C_\alpha$-$C_\alpha$ distance ≤ 15 Å extracted from the 20 best structures obtained for both states. The red circles (green stars) correspond to ECs only satisfied in ground/open (respectively excited/closed) ATP-lid state.

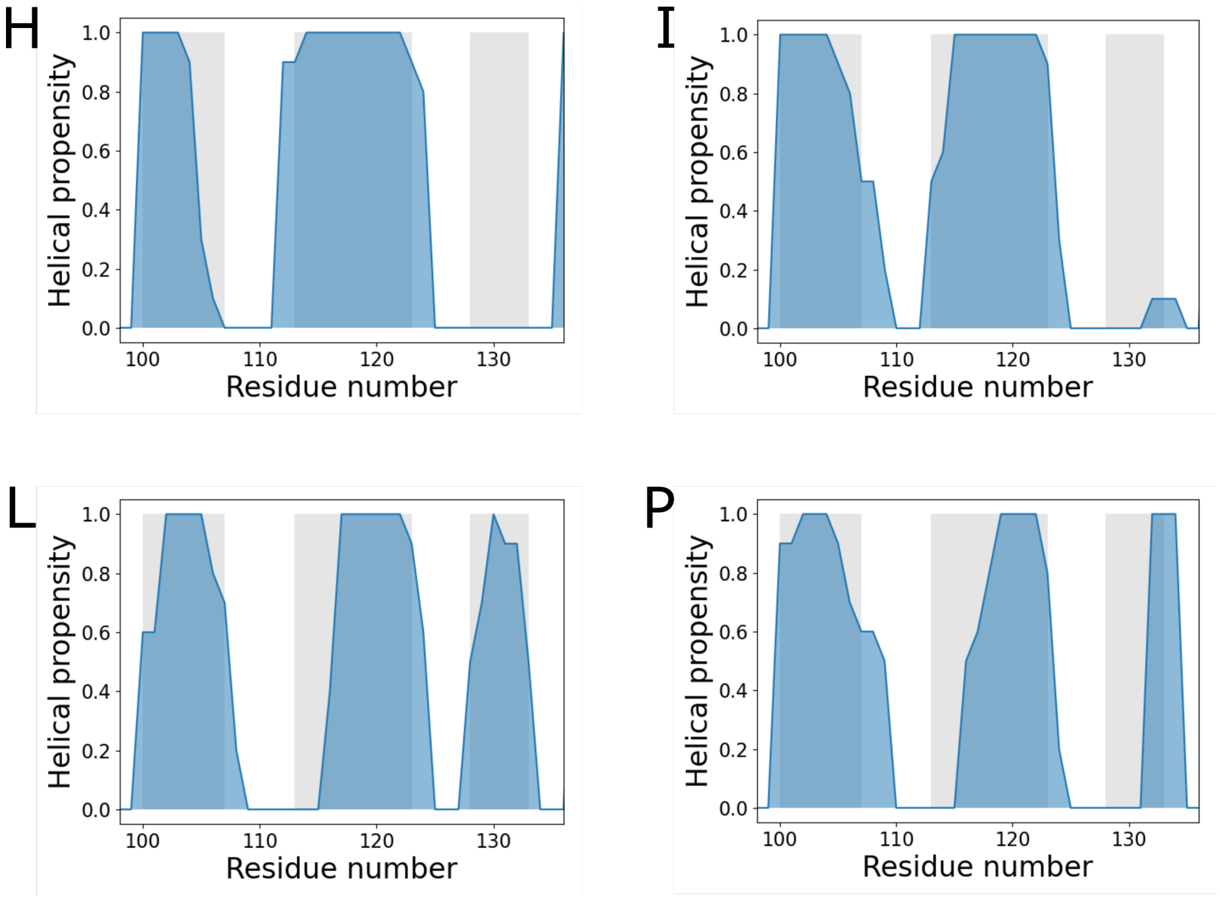


**Figure S9: Probability of helical structures along the sequence of the ATP lid, in 4 MD simulations depicting a transition of the ATP-lid from a closed to an open state.** The helical propensity are shown in blue. For comparison, the helical regions of the ground state are represented in gray (helices 3, 4 and 5). Each panel is annotated with the letter corresponding to panel of Fig. S6, presenting the 20 different MD simulations starting from closed-state conformers. These four trajectories were selected on the basis of their violations of the specific NOE of the open state, which were lower than *ca*. 3.5 Å. On average over these four selected simulations, the helical propensity of the helix-5 (residues[128-134]) is 28 ± 18%. In comparison, on average over the 20 simulations on the ground state, the helical propensity of the helix-5 is much higher: 63 ± 5% on the last 120 ns. Therefore, the formation of the helix-5 is not systematically observed after a swap of the ATP-lip from the closed to the open state.
